# Massively parallel phenotyping of variant impact in cancer with Perturb-seq reveals a shift in the spectrum of cell states induced by somatic mutations

**DOI:** 10.1101/2020.11.16.383307

**Authors:** Oana Ursu, James T. Neal, Emily Shea, Pratiksha I. Thakore, Livnat Jerby-Arnon, Lan Nguyen, Danielle Dionne, Celeste Diaz, Julia Bauman, Mariam Mounir Mosaad, Christian Fagre, Andrew O. Giacomelli, Seav Huong Ly, Orit Rozenblatt-Rosen, William C. Hahn, Andrew J. Aguirre, Alice H. Berger, Aviv Regev, Jesse S. Boehm

**Affiliations:** Broad Institute of Harvard and MIT, Cambridge, MA, USA; Howard Hughes Medical Institute; Department of Medical Oncology, Dana-Farber Cancer Institute, Boston, MA, USA; Department of Medicine, Brigham and Women’s Hospital and Harvard Medical School, Boston, MA, USA; Human Biology Division, Fred Hutchinson Cancer Research Center, Seattle, WA 98109, USA

## Abstract

Genome sequencing studies have identified millions of somatic variants in cancer, but their phenotypic impact remains challenging to predict. Current experimental approaches to distinguish between functionally impactful and neutral variants require customized phenotypic assays that often report on average effects, and are not easily scaled. Here, we develop a generalizable, high-dimensional, and scalable approach to functionally assess variant impact in single cells by pooled Perturb-seq. Specifically, we assessed the impact of 200 TP53 and KRAS variants in >300,000 single lung cancer cells, and used the profiles to categorize variants into phenotypic subsets to distinguish gain-of-function, loss-of-function and dominant negative variants, which we validated by comparison to orthogonal assays. Surprisingly, KRAS variants did not merely fit into discrete functional categories, but rather spanned a continuum of gain-of-function phenotypes driven by quantitative shifts in cell composition at the single cell level. We further discovered novel gain-of-function KRAS variants whose impact could not have been predicted solely by their occurrence in patient samples. Our work provides a scalable, gene-agnostic method for coding variant impact phenotyping, which can be applied in cancer and other diseases driven by somatic or germline coding mutations.

## INTRODUCTION

Precision medicine requires the ability to predict how specific genetic variants function in each patient (Rehm and Fowler, 2019). In cancer, one useful proxy to detect functionally selected variants is by their occurrence within patient cohorts (*e.g.*, KRAS G12V/D, TP53 R175H, BRAF V600E). However, most coding variants detected by cancer genome sequencing are rare, even within established cancer genes (Bailey et al., 2018; Lawrence et al., 2014; Tate et al., 2019; Zehir et al., 2017). Even in the case of highly recurrent variants, their mechanistic effect on cancer phenotype(s) is often undefined. As a result, distinguishing all variants that result in phenotypic changes from those that have no discernible effect remains a challenging problem that limits the interpretation of tumor genome sequencing.

Previous studies have used both computational and experimental assays to determine the putative functional impact of variants, defined as a significant difference between a variant and the wildtype allele. However, each approach has substantial limitations. Computationally, recurrent mutations that are spatially localized (Chang et al., 2016; Kamburov et al., 2015) or evolutionarily conserved (Figliuzzi et al., 2016; Hopf et al., 2017) are less likely to be functionally neutral. However, detecting recurrence while correcting for the non-random imprint of wide ranging mutagenic processes, which are only partly known (Giacomelli et al., 2018), requires substantial data sets to achieve statistical power (Alexandrov et al., 2013, 2015; Lawrence et al., 2013). Moreover, inference of positive selection on a given variant does not provide information about the specific biological function(s) it affects. Experimentally, gene-by-gene functional genomics approaches for variant impact phenotyping have assessed large numbers of alleles within a single gene, such as ERK1/ERK2 (Brenan et al., 2016), BRCA1 (Findlay et al., 2018), PI3K (Dogruluk et al., 2015; Yu et al., 2020), MEK1/2 (Gao et al., 2018) and TP53 (Boettcher et al., 2019; Giacomelli et al., 2018; Kotler et al., 2018). However, these have typically focused on specific signaling functions, and thus rely on gene- and lab-specific bespoke cellular assays, with specialized phenotypes. Such assays have limited generalizability and reproducibility, require some prior knowledge of the gene’s function, and often distinguish variants only by one dimension, without extensive information on their molecular functions.

By contrast, gene expression profiles (Berger et al., 2016; Dogruluk et al., 2015; Kim et al., 2016; Yu et al., 2020) and multi-parameter cellular imaging (Rohban et al., 2017) provide generalized phenotypes, are theoretically applicable to any gene, and yield high-dimensional phenotypes that are readily interpretable. However, to date, such approaches required arrayed, one-by-one measurements of variant impact, and were thus limited in scale. Moreover, bulk profiling could not distinguish between two types of variant impact – a uniform (or unimodal) effect across the cells, or diverse effects – multi-modal or otherwise.

Here we modified Perturb-Seq for pooled genetic screens with single cell RNA-Seq readout (Adamson et al., 2016; Dixit et al., 2016) to phenotype coding variants in a highly scalable approach for single-cell Expression-based Variant Impact Phenotyping (sc-eVIP). We benchmarked this approach by studying 200 variants of the TP53 and KRAS genes in 300,000 single cells, introduced computational methods for distinguishing the functional impact of specific variants, and demonstrated how population-based measurements may fail to capture the impact of variants on single cell heterogeneity.

## RESULTS

### A Perturb-Seq assay for coding variant phenotyping

To assess the impact of coding variants at scale, we modified Perturb-Seq to simultaneously resolve the identity of an exogenously introduced cancer variant tagged with a DNA barcode together with the induced expression state at the single cell level (Fig. 1a**, Methods**). Building on our prior work (Berger et al., 2016), we reasoned that variant function could be assessed by comparing gene expression in cells with each of the tested variant constructs to that of cells with the WT gene construct (Fig. 1a). We annotated variants as neutral if they were indistinguishable from the WT construct, and putatively impactful if they deviated significantly. To do this in a pooled setting, we cloned the coding sequence of each variant tested, each tagged with a distinct 10bp-long DNA barcode, into a modified Perturb-seq vector (**Methods**), and recovered both the expression profile of each cell and the identity of the variant(s) it overexpresses by 3’ scRNA-seq (Fig. 1a).

**Figure 1.**
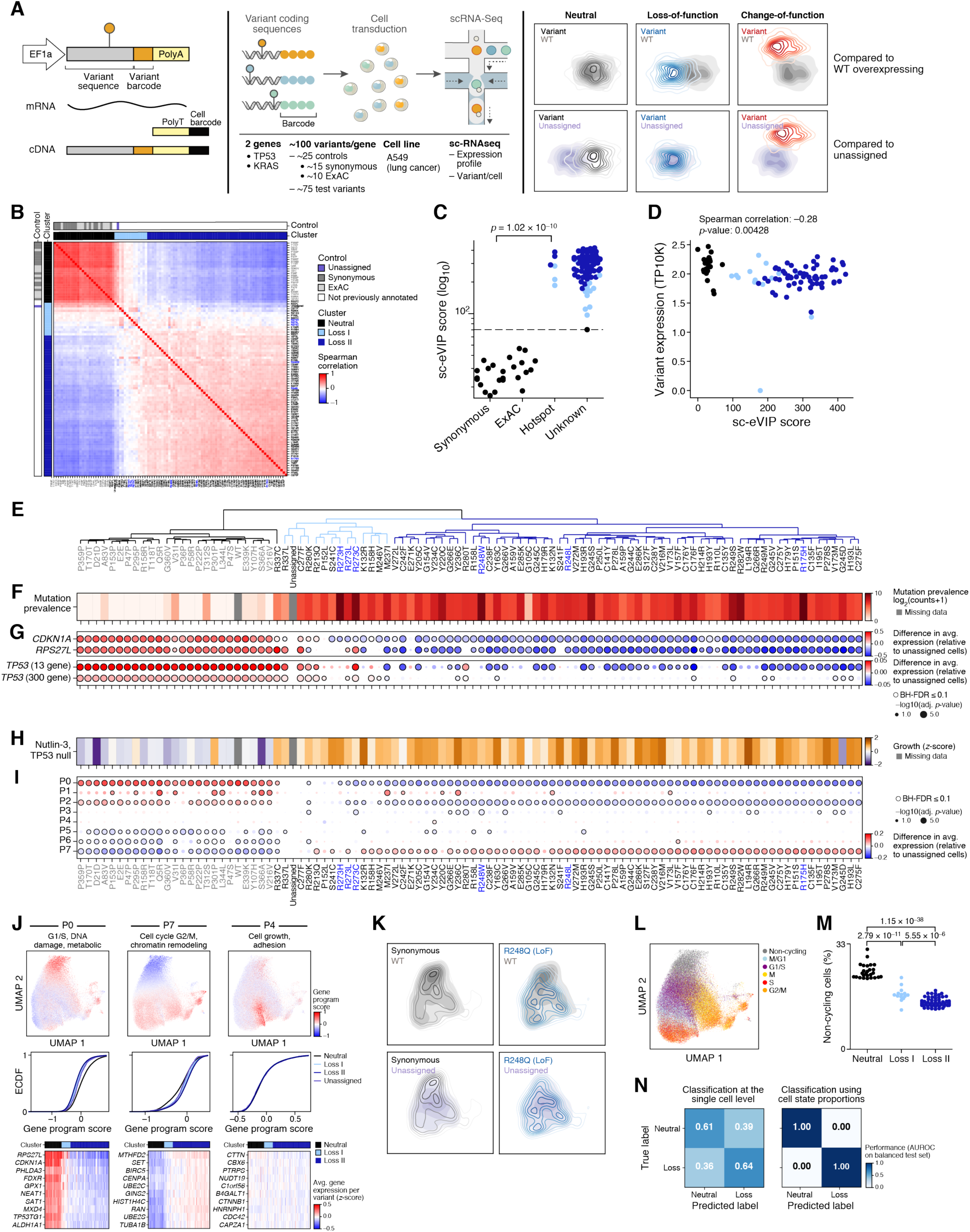
Perturb-Seq for scalable high-content coding variant profiling recapitulates known biology of loss of function variants in TP53. **a.** sc-eVIP for measuring the impact of coding variants using Perturb-seq. Left: ORF library with distinct barcodes associated with each different variant to test. Middle: transduction and Perturb-seq. Right: variant impact assessment by the deviation of profiles from cell carrying variant from those of cells overexpressing the wildtype version. **b-e.** Distinction of neutral from loss of function variants by Perturb-Seq. **b.** Spearman correlation coefficients (red/blue) between the mean profiles of each pair of variants (rows, columns), clustered into classes (black: neutral, light blue: Loss I, dark blue: Loss II, horizontal and vertical bars), and labeled by controls (dark gray: synonymous, light gray: ExAC missense variants, purple: unassigned). **c.** sc-eVIP scores (*y* axis) for variants (dots) in each category (*x* axis). Dotted line: 1% FDR. **d.** sc-eVIP scores are independent of variant expression. Variant expression (*y* axis, transcripts per 10,000 UMIs/cell (TP10K)) for variant (dots) with different sc-eVIP scores (*x* axis). **e-i.** Variant classes are associated with distinct mutation frequency, TP53 expression signatures, functional assays (growth upon treatment with Nutlin-3 in a TP53-null background) and expression programs. **e.** Hierarchical clustering of variants by the correlation profiles in **b.** Black: neutral; light blue: Loss I; dark blue: Loss II. Grey font: controls (synonymous and ExAC), blue font: hotspot variants (positions 175, 248, 273). **f.** Mutation frequency (log_2_(counts+1) of variant occurrences in a pan-cancer curated set) of each variant, ordered as in **e**. **g.** Difference (dot color) in mean expression or signature score between a variant (columns, ordered as in **e**) and unassigned cells and the significance of this difference (-log_10_(adj. p-value), Kolmogorov-Smirnov test, dot size, **Methods**) for each of two genes canonically induced by TP53 and two TP53-associated signatures (rows). Colored border: BH FDR<10%. **h.** Growth with Nutlin-3 in a TP53-null background (z-score) for each variant (ordered as in **e**). **i.** Difference (dot color) in mean program score between a variant (columns, ordered as in **e**) and unassigned cells and the significance of this difference (-log_10_(adj. p-value, Kolmogorov-Smirnov test, dot size, **Methods**) for each gene program (rows), as defined by clustering genes (**Methods**). Program 1, higher in assigned *vs*. unassigned cells was enriched for translation, nonsense-mediated decay, and viral transcription, and may reflect the response to lentiviral transduction. Colored border: BH FDR<10%. **j.** Gene programs vary across variant classes. Top: UMAP embedding of single cell profiles, colored by program scores (color bar). Middle: Cumulative distribution function of the program scores (*x* axis) for each variant class (color). Bottom: average expression (z-score, color bar) in cells of each variant (columns) of genes (rows) most correlated with the mean of the expression program. **k.** Variant induced shift in cell distributions. Density map of cell profiles organized in a 2-dimensional UMAP embedding, comparing the density of cells overexpressing a synonymous allele (black, right) or a loss of function variant, R248W (blue, left) to either the WT TP53 allele (grey, top) or unassigned cells (purple, bottom). **l,m.** Reduced proportion of non-cycling cells in loss of function TP53 variants. **l.** UMAP embedding of single cell profiles, colored by their assignment to cell cycle phases. **m.** Proportion of non-cycling cells (*y* axis) among cells carrying each variant (dots) across variant classes (*x* axis). Adj. p-value: t-test. **n.** Accurate variant classification by mean profiles but not at the single cell level. Performance (AUROC on balanced test set, color bar) of logistic regression classifiers predicting the class of variant for individual cells (left), or for the entire set of cells (by proportion of cell states) (right).

As a first test case, we assessed 75 cancer-associated coding variants in TP53, compared to synonymous controls and non-synonymous common variants. To this end, we selected and synthesized 100 TP53 variants (99 passed QC; Extended Data Fig. 1a, **Extended Data Table 1**). These included (**1**) the 75 most recurrent TP53 mutations from TCGA (Bailey et al., 2018), MSKCC-IMPACT (Zehir et al., 2017), and GENIE (AACR Project GENIE Consortium, 2017), which are predicted to be uniformly loss-of-function (Tate et al., 2019); (**2**) 15 synonymous variants as controls (expected to be indistinguishable from the wildtype allele); and (**3**) 10 non-synonymous variants from healthy cohorts (ExAC, (Lek et al., 2016)). We transduced the 99 pooled TP53 variants into A549 lung cancer cells, a known suitable biosensor for TP53 function (Giacomelli et al., 2018) at low MOI to favor single variants per cell (estimated MOI 0.77 and detection probability 0.78). We selected for successfully infected cells, and performed scRNA-seq (Fig. 1a). Because the expression level of the variants was sufficiently high, we did not perform a dial-out PCR (Dixit et al., 2016) to enrich variant barcodes.

Overall, we associated variants and profiles reliably. Specifically, we recovered 162,314 high quality cells of which 84% had detectable variant barcodes and 62% were confidently annotated with a single expressed variant (median 926 high confidence cells per variant; Extended Data Fig. 1b,c). In >70% of cases, a variant overexpressed in a cell was supported by at least 2 barcode UMIs (Extended Data Fig. 1d). Furthermore, each of the TP53 variants was expressed at levels comparable to the wild type construct with only two variants exceeding a 1.5-fold expression difference (Fig. 1d, Extended Data Fig. 1e,f, M237I and Y236C). To reduce potential biases due to variant overexpression levels, we regressed out the variant barcode expression in each cell.

### Single-cell expression variant impact scores correctly distinguish TP53 loss-of-function variants

To score and categorize variants by their expression profiles we used two complementary approaches: scoring the distance between the mean profiles of variants and the WT construct by extending our previous expression-based variant impact phenotyping (eVIP) approach (Berger et al., 2016), and unsupervised clustering of variants by their mean profiles. Our “single cell eVIP” scores (sc-eVIP) quantify the extent to which cells overexpressing a variant deviate from the mean expression profile of cells overexpressing the wildtype allele (**Methods**) using Hotelling’s T^2^ test (Hotelling, 1931), a multivariate generalization of a t-test applied to a low-dimensional representation of cells in principal component (PC) space (**Methods**). This statistical test yielded highly concordant results, even when we down-sampled the cells ∼10-fold to 100 cells/variant; (Extended Data Fig. 1i). We then called impactful variants at a 1% FDR by comparison to an empirical null distribution generated from comparisons between control synonymous variants (**Methods**). As a complementary unsupervised approach, we hierarchically clustered the variants using an *L*1 distance and complete linkage, applied to the correlation matrix of average expression profiles (**Methods,** Fig. 1b,e). Finally, we investigated the gene programs underlying the different classes of variants tested, by clustering the average gene expression profiles across variants to identify sets of genes with similar behaviors across variants (Extended Data Fig. 2f, **Methods**) or by principal component analysis (**Methods**, Extended Data Fig. 2g-k).

**Figure 2.**
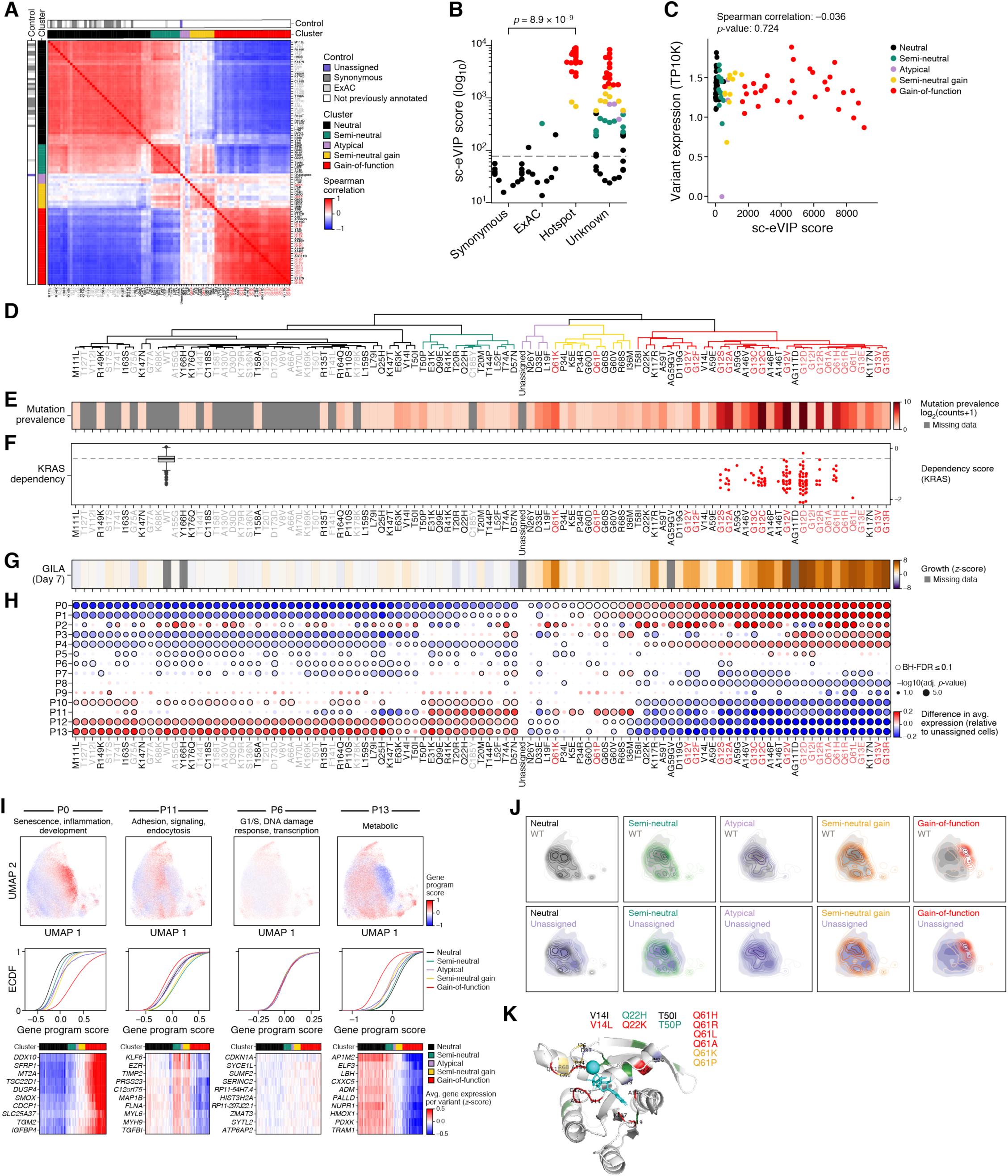
sc-eVIP correctly annotates known gain-of-function variants in the KRAS oncogene and reveals five functional classes of KRAS coding mutations. **a-c.** Distinction of neutral from impactful variants in KRAS by Perturb-Seq. **a.** Spearman correlation coefficients (red/blue) between the mean profiles of each pair of variants (rows, columns), clustered into classes (black: neutral, green: semi-neutral, purple: atypical, gold: semi-neutral gain, red: gain-of-function, horizontal and vertical bars), and labeled by controls (dark gray: synonymous, light gray: ExAC missense variants, purple: unassigned). **b.** sc-eVIP scores (*y* axis) for variants (dots) in each category (*x* axis). Dashed line: 1% FDR. **c.** sc-eVIP scores are independent of variant expression. Variant expression (*y* axis, transcripts per 10,000 UMIs/cell (TP10K)) for variant (dots) with different sc-eVIP scores (*x* axis). **d-h.** Variant classes are associated with distinct mutation frequency, KRAS dependency, growth in low attachment (GILA) phenotypes, and expression programs. **d.** Hierarchical clustering of variants by the correlation profiles in **a.** Grey font: controls (synonymous and ExAC), red font: hotspot variants (positions 12, 13 and 61). **e.** Mutation frequency (log_2_(counts+1) of variant occurrences in a pan-cancer curated set, color bar) of each variant, ordered as in **e**. **f.** Dependence of cell line growth on KRAS (*y* axis), for cell lines (dots) categorized by their KRAS genotype status (*x* axis). Gray: wildtype KRAS, red: missense KRAS variants. **g.** Growth in low attachment of HA1E cells (z-score, color bar) for each variant (columns, ordered as in **e**). **h.** Difference (dot color) in mean expression or signature score between a variant (columns, ordered as in **e**) and unassigned cells and the significance of this difference (-log_10_(adj. p-value, Kolmogorov-Smirnov test, dot size, **Methods**) for each gene program (rows), as defined by clustering genes (**Methods**). Colored border: BH FDR<10%. Program 2, higher in assigned vs. unassigned cells was enriched for translation, nonsense-mediated decay, viral processes and metabolism, and may reflect the response to lentiviral transduction. **i.** Gene programs varying across variant classes. Top: UMAP embedding of single cell profiles (dots), colored by program scores (color bar). Middle: Cumulative distribution function (CDF) of program scores (*x* axis) for each variant class (color). Bottom: mean expression (z-score, color bar) of genes (rows) most correlated with the mean of the expression program in cells of each variant (columns). **j.** Variant-induced shift in cell distributions. Density map of cell profiles organized in a UMAP embedding, showing the density of cells overexpressing each variant class (colored as in **a**) and either the WT KRAS allele (grey, top) or unassigned cells (purple, bottom). **k.** Annotation of variant classes on the 3-dimensional structure of the KRAS protein. Each position is colored by the variant class with the highest impact assigned to variants at that position, with 4 variants assigned to multiple categories highlighted (listing all variants at the position, each colored by its assigned class, as in **a**)

Both the sc-eVIP scores and unsupervised clustering correctly distinguished expected loss-of-function variants from the WT and control variants. Specifically, all (25/25; 100%) synonymous and ExAC control variants exhibited sc-eVIP scores similar to WT (FDR 1%, Fig. 1c), and formed a separate cluster, together with R337C (Fig. 1e, black). The remaining 73/74 variants (98.6%), including those at hotspot positions 175, 248 and 273 (top 3 most frequent variants in COSMIC (Tate et al., 2019), Fig. 1e, blue) exhibited significant sc-eVIP scores and formed a distinct cluster from the neutral controls (Fig 1. b,e). This cluster included the average profile from all unassigned cells (cells without a detected variant barcode), suggesting the variants in this cluster were likely loss-of-function. The 25 control TP53 variants significantly induced canonical signatures of TP53 overexpression compared to unassigned cells (Fischer, 2017; Jeay et al., 2015), including induction of *CDKN1A* and *RPS27L* expression (Fig. 1g), and showed the expected increases in the proportion of non-cycling cells, consistent with cell cycle arrest, as assessed from their single-cell expression signatures (Fig. 1m, Extended Data Fig. 3g).

**Figure 3.**
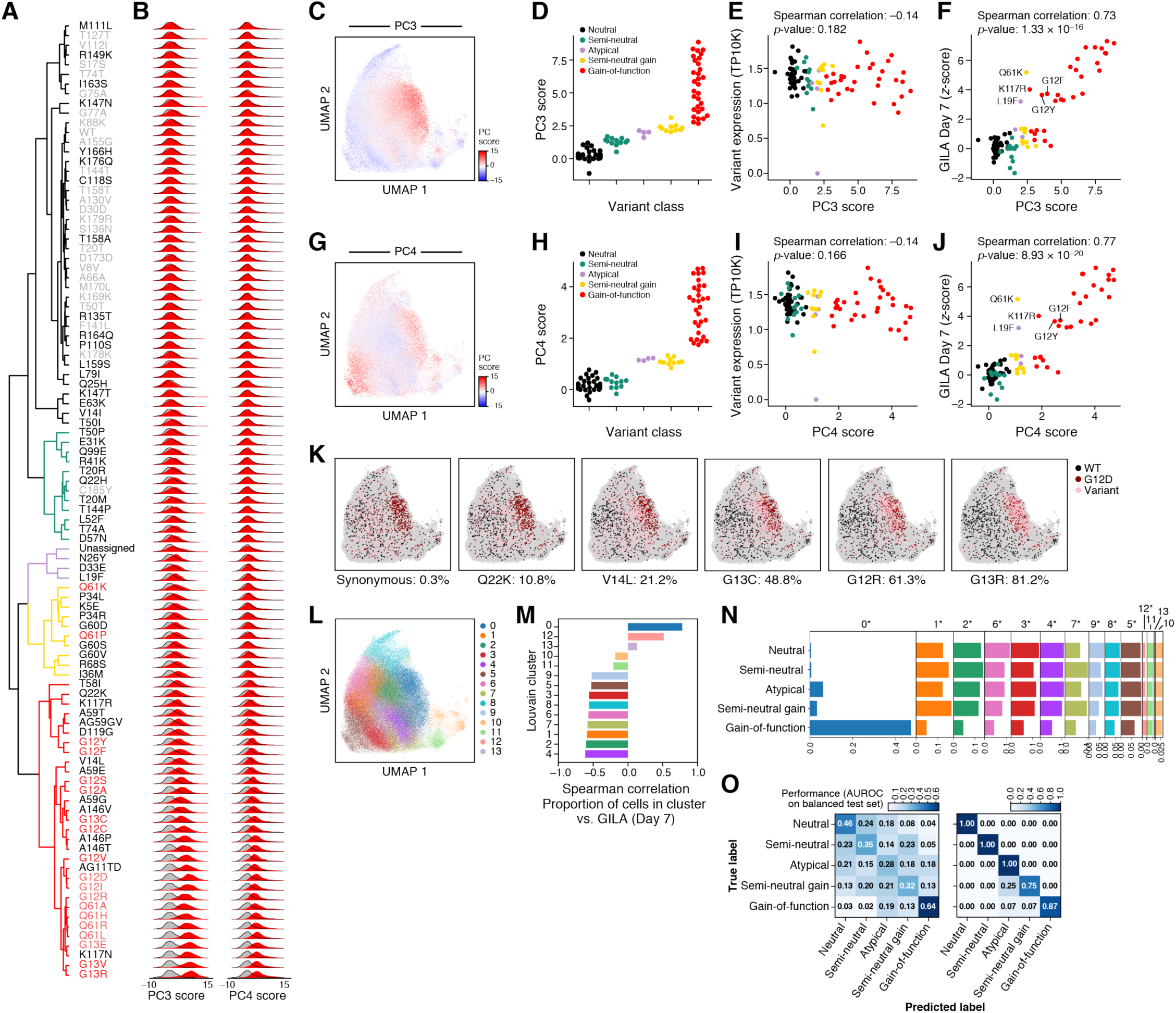
KRAS variants form a gradual functional gradient within and across variant groups. **a-b.** Continuous variation across KRAS variants. **a**. Hierarchical clustering of variants by the correlation profiles in Fig 2a. black: neutral, green: semi-neutral, purple: atypical, gold: semi-neutral gain, red: gain-of-function. Grey font: controls (synonymous and ExAC), red font: hotspot variants (positions 12, 13 and 61). **b.** Distribution of the scores of principal components (PCs) 3 and 4 for cells carrying each variant (red) and WT KRAS overexpressing cells (gray). **c-j.** PC 3 and 4 scores are concordant with functional assays and independent of variant overexpression levels. **c,g.** UMAP embedding of single cell profiles (dots), colored by PC scores (color bar). **d,h**. Mean PC scores (*y* axis) for each variant (dots), from the five variant classes (*x* axis, colored as in **a**). **e.i**. Normalized variant barcode expression level (*y* axis, transcripts per 10,000 UMIs/cell (TP10K)) and sc-eVIP impact scores (*x* axis) for each variant (dots), colored by variant class. **f,j**. GILA scores (*y* axis) and mean PC scores (*x* axis) across variants (dots), colored by variant class. **k-n**. Variation in cell state proportions across variants. **k**. UMAP embedding of single cell profiles (dots), colored by WT KRAS (black), gain-of-function G12D (dark red), each of 5 variants (pink, label at bottom), and all other cells (grey). Bottom: fraction of cells of the noted variant present in gain-of-function-associated cell state 0 (as in **l**) for each variant. **l**. UMAP embedding as in **k**, colored by cell clusters. **m**. Spearman correlation coefficient (*x* axis) for each cluster in **l** (*y* axis) between the proportion of cells in each cluster in **l** (*y* axis) and the functional assay (GILA). **n**. Fraction of cells in each cell cluster (*x* axis) from each variant class (*y* axis). **o**. Accurate variant classification by mean profiles but not at the single cell level. Performance (AUROC on balanced test set, color bar) of logistic regression classifiers predicting the class of variant for individual cells (left), or for the entire set of cells (by mean expression profile) (right).

In this cellular context, the variants that were not in the neutral cluster further partitioned into two groups (Loss I and Loss II) by both increasing sc-eVIP scores (Fig. 1c) and as sub-clusters (Fig. 1e, light and dark blue). The groups varied in the strength of impact on canonical TP53 signatures, such that Loss I variants induced canonical TP53 signatures to a lesser degree than neutral variants, whereas Loss II variants did not affect (or repressed) TP53 signatures relative to unassigned cells (Fig. 1g, t-test between 13-gene and 300-gene signature scores: neutral *vs* Loss I p=2.9*10^-15^, and 7.4*10^-17^, and Loss I vs Loss II: 1.4*10^-10^ and 2.7*10^-10^). The repression of canonical TP53-induced genes CDKN1A and RPS27L, and to a lower degree TP53 signatures in a majority of Loss I and II variants is consistent with the annotation of 71/74 of these variants as having dominant negative effects in previous work (Giacomelli et al., 2018) (all Loss I and II variants except R337L, R280K and G105C, as defined by a functional assay of growth upon treatment with Nutlin-3 in a TP53-WT background, z-score higher than 0.61(Giacomelli et al., 2018)). Specifically, 68/71 and 62/71 of dominant negative variants show significant repression of CDKN1A and RPS27L and 40/71 and 10/71 for 13-gene and 300-gene TP53 signatures respectively.

As expected, loss of function and neutral variants had diametrically-opposed effects on multiple programs (**Methods**) compared to unassigned cells: program 0 (G1/S checkpoint, DNA damage response and metabolism) and 2 (adhesion, differentiation, migration) were repressed by loss of function variants and activated by neutral variants (Fig. 1i,j, Extended Data Fig. 2a-e), whereas program 7 (cell cycle G2/M, chromatin remodeling) was activated by loss-of-function variants and repressed by neutral ones (Fig. 1i,j, Extended Data Fig. 2a-e).

Our sc-eVIP scores and the associated gene programs from Perturb-Seq were highly concordant with those from an optimized TP53-specific cellular assay for growth under Nutlin-3 treatment, in a TP53-null background which was previously conducted with the same cell line and over-expression constructs (Giacomelli et al., 2018) (Spearman ρ=0.73, p= 3.2*10^-17^, Fig. 1h, Extended Data Fig. 3a), as well as additional assays (Extended Data Fig. 3a-f)

At the single cell level, neutral and loss-of-function variants did not occupy mutually exclusive cell states (Fig. 1k, Extended Data Fig. 4g), and the two groups showed extensive cell state overlap, but differed in their distribution across these states, especially in cell cycle phase distribution (Fig. 1k,l,m, Extended Data Fig 3g). As a result, a logistic regression classifier using gene expression to predict if any individual cell harbors a loss-of-function or neutral variant had limited accuracy (area under the precision-recall curve (AUPRC) = 0.78 on a test set of 50% of variants) (Fig. 1n), but overall variant classification using the proportions of cells in each of 15 cell subsets (as defined by Louvain clustering) was highly accurate (AUPRC=1), suggesting that our TP53 variant phenotypes mostly reflect a shift in cell state distributions (Fig. 1n).

**Figure 4.**
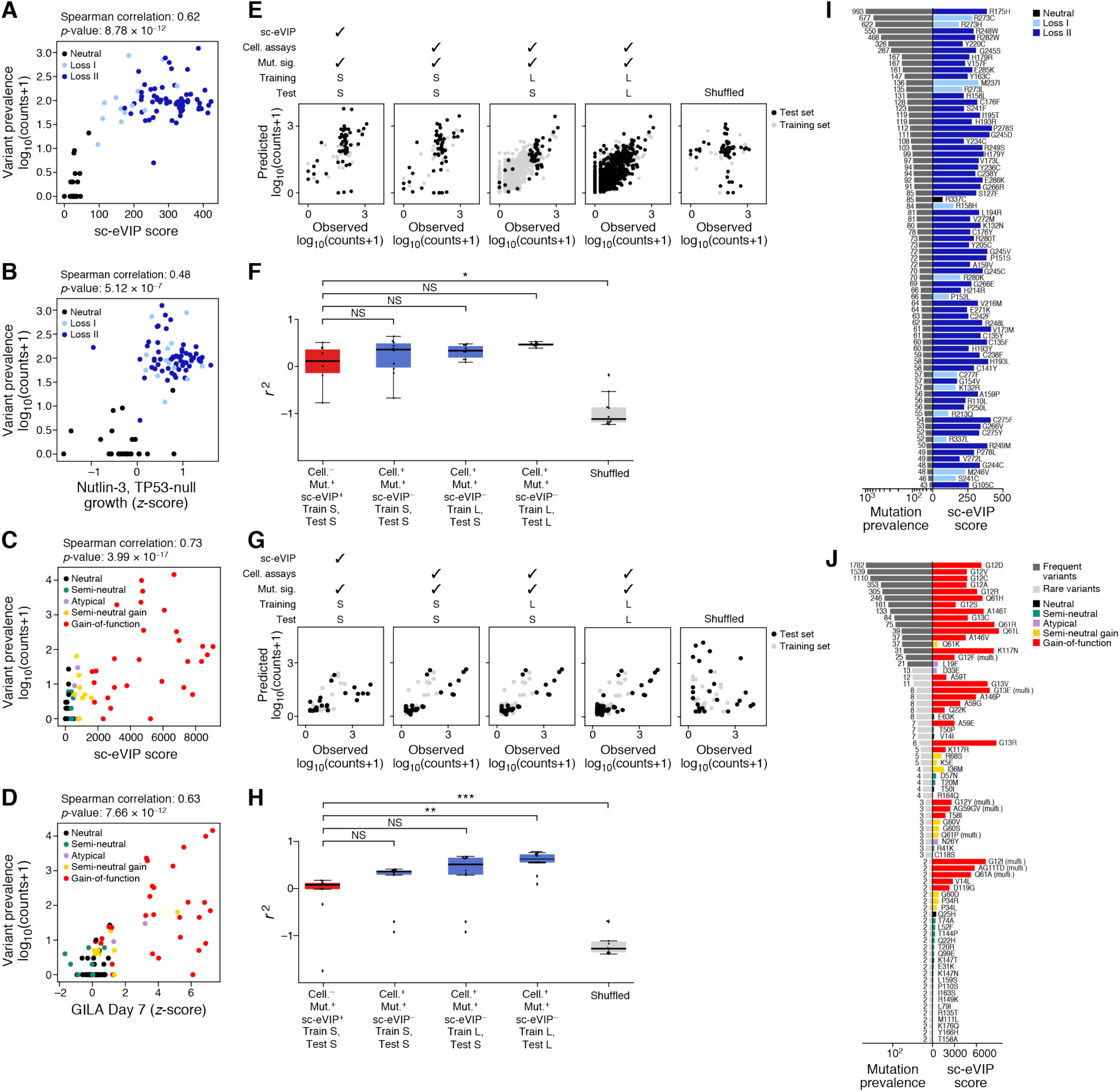
Relationship between sc-eVIP scores and the frequency of variants in patient cohorts. **a-d.** Comparison of sc-eVIP scores and functional assays with variant prevalence in cancer cohorts for TP53 (**a,b**) and KRAS (**c,d**). **a,c**. Mutation prevalence (*y* axis, log_10_(counts of mutation in cohort + 1) and sc-eVIP scores (*x* axis) across TP53 (a) and KRAS (b) variants (dots), colored by variant class. **b,d**. Mutation prevalence (*y* axis, log_10_(counts of mutation in cohort + 1) and functional assay scores for TP53 (**b**, growth with Nutlin-3, in a TP53-null background, z-score, *x* axis) and KRAS (**d**, GILA, z-score, *x* axis) across variants (dots), colored by variant class. **e-h**. Comparison of generalized linear models for predicting variant prevalence in cancer cohorts using mutational signatures and either sc-eVIP or functional assays for TP53 and KRAS. **e,g**. Top: Specification of 5 compared models, trained on the subset of variants profiled in this study (small set, S) or on a large dataset of thousands of variants (L) or by shuffling the observed variant prevalence across variants (**Methods**). Observed (*x* axis) and predicted (*y* axis) variant prevalence (log_10_(counts + 1)), for each model across variants (dots), colored by whether the variant is in the training (gray) or test (black) set, for TP53 (**e**) and KRAS (**g**). **f,h**. Coefficient of determination (r_2_) of each of the 5 models for TP53 (**f**) and KRAS (**h**) relative to a model predicting the mean variant prevalence, across 10 models, colored by whether they use sc-eVIP scores for prediction (red), functional assays (blue), or neither (gray). **i,j**. Mutation prevalence (*x* axis, left, colored by mutation prevalence (dark gray >= 20 counts and light gray <20 counts) and sc-eVIP scores (x axis, right, colored by variant class), for each of the TP53 (**i**) and KRAS (**j**) variants in the study.

### Perturb-Seq for KRAS coding variants identifies rare, gain-of-function mutations and annotates additional KRAS mutations

Given the recent success in therapeutically targeting specific KRAS variants (Hong et al., 2020), we next evaluated the utility of coding variants Perturb-seq for a series of 98 KRAS variants. Although A549 cells harbor a KRAS G12S allele, we previously found that expression of KRAS alleles in this cell line permits the discrimination of the function of exogenously expressed KRAS alleles in functional assays (Berger et al., 2016; Kim et al., 2016; Singh et al., 2009).

We selected the 75 most recurrent KRAS alleles in cancer cohorts and 26 negative control alleles, including 16 synonymous variants and 10 common, non-synonymous variants from ExAC (98 passed QC; Extended Data Fig. 5a, **Extended Data Table 2**). These alleles included those reported frequently in cancers (n=1,782, 1,539, 1,110 for G12D, V, C, respectively) as well as 34 rare alleles observed in fewer than 5 individuals among TCGA (Bailey et al., 2018), MSKCC-IMPACT (Zehir et al., 2017), and GENIE (AACR Project GENIE Consortium, 2017) databases. We analyzed 150,044 high-quality single cell profiles (68% annotated to a single variant, a median of 1,058 cells per variant, Extended Data Fig. 5b,c,d, e). As for TP53, most variant constructs were expressed at similar levels (Extended Data Fig. 5f), with only 2 outlier constructs with reduced expression, both at position 61 (Q61K and Q61L, respectively), and no apparent relationship between variant expression levels and sc-eVIP scores (Fig. 2c, Spearman ρ=-0.04, p=0.72).

**Figure 5.**
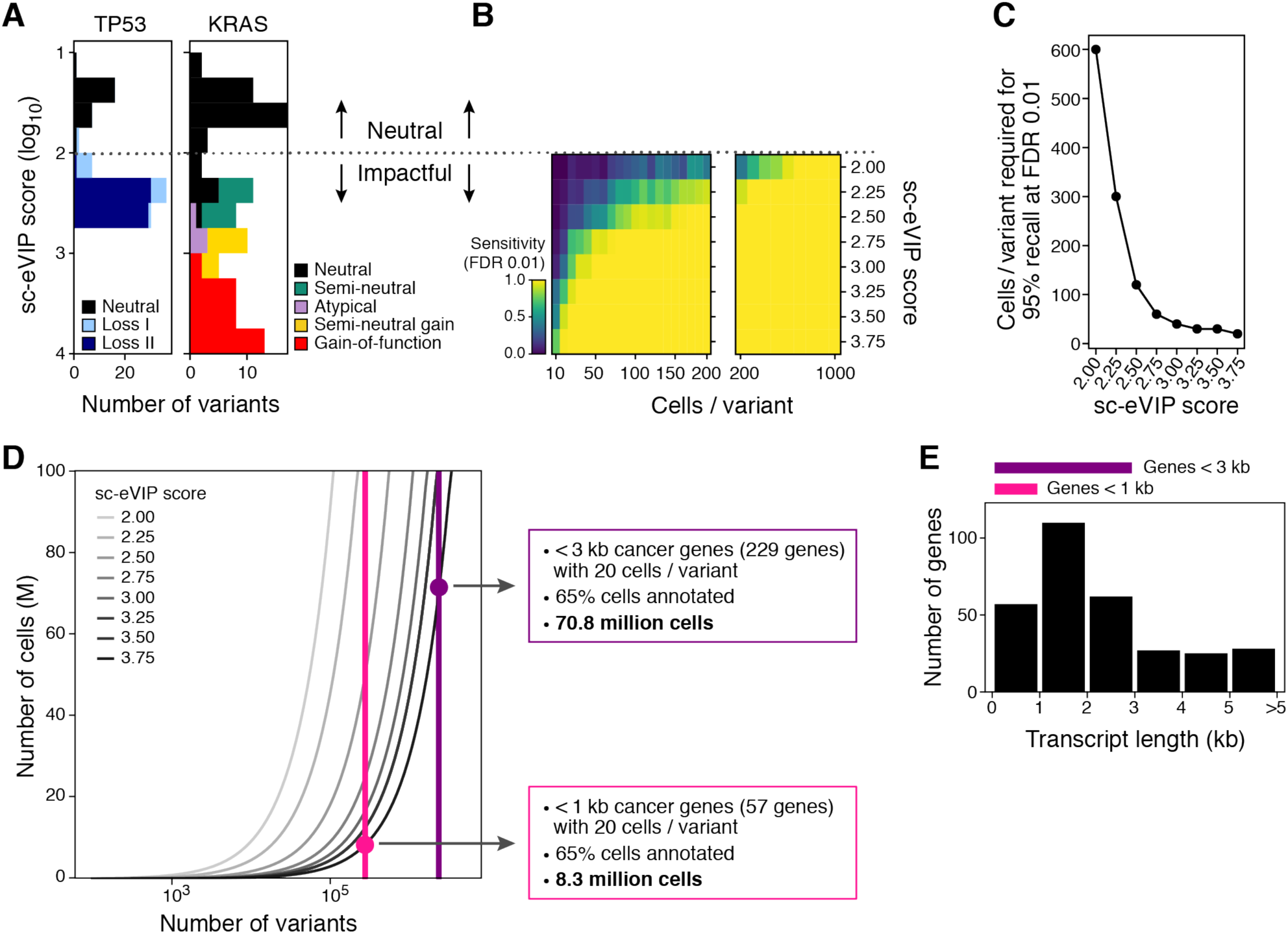
Power analysis and outlook for the variant-to-function efforts to phenotype cancer coding variation. **a-c.** Power analysis for detecting impactful variants as a function of effect size. **a**. Variant impact effect size (*y* axis, log_10_(sc-eVIP score)), colored by variant class for TP53 (left) and KRAS (right) variants. Dotted line separates impactful from neutral variants. **b**. Sensitivity (colorbar) of impactful variant detection for each variant effect size bin in **a** (rows) at increasing numbers of cells profiled per variant (columns). **c.** Number of cells required for a sensitivity of 0.95 at an FDR of 1% (*y* axis) for each variant effect size bin (*x* axis). **d,e.** Projected number of cells required for a cancer coding variant impact atlas. **d**. Number of cells (millions, y axis) required to profile for testing a given number of variants (*x* axis), at different variant impact effect sizes (colored curves). Vertical lines: number of variants needed for studying all cancer genes in the Foundation Medicine Panel less than 1kb in length (magenta) or less than 3kb (purple). Dots: number of cells required for these atlases for detection at the strongest effect size bin. **e**. Distribution of transcript lengths (*x* axis, Kb) for the 309 cancer genes in the Foundation Medicine panel.

Both impact scores (Fig. 2b, Extended Data Fig. 5i) and clustering by mean expression profiles (Fig. 2a,d) correctly distinguished control synonymous KRAS alleles from known gain-of-function variants at hotspot positions 12, 13 and 61 (Fig. 2b, P<8.9*10^-9^, t-test). Based on both scores and clusters, we annotated another 19 variants as neutral, including 9 of 10 ExAC control variants and 10 variants observed in patients at low frequency (Y166H, T58A, C118S, K176Q, R135T, R164Q, L79I, R149K and I63S; each in fewer than 4 patients, Fig. 2e). Previously well-characterized gain-of-function variants at positions 12, 13 and 61 had higher sc-eVIP scores than neutral alleles and separated in distinct clusters (Fig. 2a,b,d). Moreover, in evaluating data from the Broad Cancer Dependency Map (Meyers et al., 2017), cell lines with impactful variants were more sensitive to loss of KRAS by CRISPR/Cas9 knockout than those with a wildtype allele (Fig. 2f, p<8.3*10^-147^, t-test). Finally, there was good correlation (Fig. 2g, Spearman correlation 0.75, p=1.93*10^-18^) between sc-eVIP scores and those from a phenotypic assay measuring growth in low attachment (GILA (Rotem et al., 2015), z-scores) in human embryonic kidney cells (HA1E) overexpressing KRAS alleles (Ly, 2018). Variants deemed impactful by GILA had higher sc-eVIP scores (p<3.2*10^-5^, t-test) and sc-eVIP scores were predictive of GILA high-scoring variants (z-score, AUPRC= 0.92).

### KRAS variants partition to several classes by their impact on expression

To further understand the range of KRAS variants and the gene expression programs that underlie their functional impact, we defined five variant clusters by correlation between mean single cell profiles (pseudo-bulk) (Fig. 2a,d), as well as used Louvain clustering of the pseudo-bulk expression profiles across variants to identify expression programs associated with variants independent of cluster (Fig. 2h, i, Extended Data Fig. 6a-g). In most (10 of 14) positions with more than one non-synonymous variant tested, the different non-synonymous variants were in the same cluster (Fig. 2k). One cluster captured gain-of-function variants, including those at known hotspot positions 12, 13, and 61, and had the highest sc-eVIP scores (Fig. 2a,d, red text, Extended Data Fig. 8a). A second cluster contained all synonymous (WT) variants and 9 of 10 common non-synonymous (ExAC) variants, suggesting it captured the group of neutral variants in our set (Fig 2a,d, **black**). Three remaining clusters had lower sc-eVIP scores though not as low as for the WT cluster (p<2.5*10^-10^, 9.4*10^-14^, 1.5*10^-20^ compared to WT cluster; t test), and included variants with increasing distinction by sc-eVIP scores (Fig 2a,b,d, green, purple, yellow, Extended Data Fig. 8a).

Variants in the gain-of-function cluster included positions 12, 13, 22, 59, 61, 146, 117 and 119 (Fig. 2d), which largely fall either at or near nucleotide binding domains (UniProt Consortium, 2019) (Fig. 2k, Extended Data Fig. 8f). All of these variants significantly induced program 0 (senescence, inflammation, development) (Fig. 2h,i), as well as program 1 (response to stimulus, apoptosis, secretion) and all but one induced program 4 (hypoxia, immune, stress, G2M, cell polarity) (Extended Data Fig. 6a-e). These alleles also repressed programs 10 (metabolic, G1/S, adhesion, regulation of senescence), 12 (signaling, MAPK, metabolic, development) and 13 (metabolism), and all but two additionally repressed program 11 (adhesion, signaling, endocytosis) (Fig 2h,i, Extended Data Fig. 6a-e).

The variants in the neutral cluster (Fig. 2d, black) included 18 variants observed in patients, 8 of which had low but significant sc-eVIP scores (P110S, L159S, Q25H, K147N, K147T, E63K, V14I, T50I) including variants appearing seven (V14I) or eight (E63K) times in patient cohorts (Fig. 2e). Although members of the same cluster, these variants had distinguishing features, as many repressed gene program 11 (adhesion, signaling, endocytosis) (Fig. 2h), although their impact on other programs was the same as neutral variants (Fig. 2h).

We denoted the three clusters of variants that more closely, but not precisely, phenocopied neutral as “semi-neutral” (green), “atypical” (purple) and “semi-neutral-gain” (purple) based on the gradual progression of their sc-eVIP scores from neutral to gain of function (Extended Data Fig. 8a). Both the semi-neutral and semi-neutral-gain groups included variants near nucleotide binding sites, effector binding, and the allosteric lobe (Fig. 2k, Extended Data Fig. 8f), and induced an adhesion, signaling and endocytosis program (program 11, Fig. 2h), which was lower in both gain-of-function and neutral variants. However, semi-neutral-gain variants had higher levels of gain-of-function-associated programs for senescence, inflammation, development (program 0), response to stimulus, apoptosis and secretion (program 1) and hypoxia, immune, stress, G2M, cell polarity (program 4), and lower levels of neutral-associated programs metabolic, G1/S, adhesion and regulation of senescence (program 10), signaling, MAPK, metabolic and development (program 12) and metabolism (program 13), suggesting these alleles resemble gain-of-function cell states (Fig. 2h).

Variants in the final “atypical” cluster (Fig. 2a,d, purple) showed a mixture of features from other variant groups: repression of program 0 (as in neutral and semi-neutral variants), and of programs 12 and 13 (as in gain-of-function) (Fig. 2h). Of these variants, D33E has tumorigenic potential (Kim et al., 2016), due to altered protein dynamics including changes in the switch 1 conformational state (Lu et al., 2018); and L19F conferred fitness in NIH3T3 cells (Akagi et al., 2007) and scored highly in the GILA assay.

### KRAS variants form a gradual functional gradient within and across groups

We further examined the KRAS variants that scored as impactful, leveraging the single cell profiles. We performed Principal Component Analysis (PCA) of all cells and searched for PCs that were significantly higher in cells carrying previously known gain-of-function variants from hotspot positions 12, 13 and 61 compared to cells carrying synonymous variants. Of 32 significantly different PCs (by t-test, 1% FDR), PC3 and PC4 scores (Fig. 3b) were particularly good at distinguishing between individual cells with activating *vs*. neutral variants (AUROC on balanced data 0.9 and 0.8 respectively, compared to the next best performance of 0.67 for PC5, Extended Data Fig. 6h-k). Top ranked genes for PC3 were enriched in oncogene-induced senescence, regulation of apoptosis and multiple metabolic pathways, while top genes for PC4 were enriched for adhesion.

Although PC3 scores of all neutral variants were mostly comparable to wildtype overexpressing cells (Extended Data Fig. 6j,k), other variants arranged on a continuum of increasing strength, as reflected by the increasing separation of the distribution of PC3 scores (Fig. 3c,d, Extended Data Fig. 8c), gradually shifting from the mildest separation appearing in variants in the neutral cluster that have significant sc-eVIP scores (e.g., V14I) to the strong separation in the full gain-of-function variants. Even within the gain-of-function cluster, PC3 scores varied continuously. This continuum was not explained by technical considerations, such as differences in variant overexpression levels (Fig. 3e, Spearman correlation −0.14, p=0.18) and was consistent with GILA assay (Ly, 2018) scores (Fig. 3f, Spearman correlation 0.73, p=1.3*10^-16^), for which it had high predictive power (auPRC 0.91), with only 5 of 24 variants scoring high in GILA (z-score > 3) not showing the expected increases in PC3 (Q61K, K177R, L19F, G12Y, F) (Fig. 3f, Extended Data Fig. 8c). PC4 scores varied most within the gain-of-function cluster, with neutral and semi-neutral variants showing similar levels, and atypical and semi-neutral-gain falling between neutral and gain-of-function levels. This suggests that two sources of variation may be at play, as also indicated by gene program scores (Fig. 3b,g-j, Extended Data Fig. 8c).

### KRAS gain-of-function variants largely redistribute cells within an existing phenotypic landscape

These continuous differences between variants can arise because variants lead to a new cell state, not observed in the context of WT KRAS, because of a redistribution in the same phenotypic space, or both. To distinguish these possibilities, we embedded all the single cell profiles from our experiment in two dimensions (Fig. 3l**, Methods**) and compared the distribution of single cell profiles from variants in the five categories to those from either WT or unassigned cells (Fig. 3n).

Impactful and neutral variants occupied a largely overlapping cell state space, but with a continuous shift in the distribution of cells across this space (Fig. 3k), from neutral, to semi-neutral, semi-neutral-gain, atypical then followed by the continuum of gain of function variant spectrum. Only the strongest gain-of-function variants occupied almost exclusively a cell state space (Fig. 3k,l,n, cell state “0”) barely occupied by WT-overexpressing cells (<2%), but present in some of the unassigned cells (11.7%). For example, >81% of the cells with the strongest gain-of-function variant, G13R, are in this space (Fig. 3k). This cell state is associated with high expression of gain-of-function programs 0 and 1 (Extended Data Fig. 6g). The proportion of cells in this portion of the cell state space corresponds to the observed continuum of variant activating levels by GILA (Spearman correlation = 0.78, P-value=1.8*10^-20^, Fig. 3f, Extended Data Figure 7c-d). While the gain-of-function variants also have a higher proportion of cells in M phase, and a lower proportion in S phase (Extended Data Fig. 7e), these changes are modest and, unlike in TP53, are not a major contributor to the difference between variants.

Consistent with a model of redistribution of cells across an existing state space (Fig. 3k), when we trained a multi-class logistic regression model to classify each individual cell into its corresponding variant class as defined by unsupervised clustering, we found a broad swath of misclassifications (Fig. 3o**, left**). The best performance is for gain-of-function variants, due to the high enrichment of cells in cell state 0 largely depleted of other variants (Fig. 3k-m, Extended Data Fig. 7f). Conversely, a model trained and tested on mean expression profiles has near-perfect performance, in all but two classes, suggesting that variants impact cell composition (Fig. 3o**, right**).

### Expression impact can help predict mutation frequencies, but mutation frequency alone does not always predict impact for rare KRAS variants

In principle, variants exerting stronger functional effects would be under stronger selective pressure in tumors and in many cases would be found to be mutated at higher frequencies across patients. Thus, we related our functional characterization of variants to observed mutation frequencies in cancer cohorts, to test if functional effects can help predict mutation frequency, and conversely, to understand the degree to which variant impact in our assay can be ascertained based on mutation frequency alone. There were significant correlations between the number of times a variant occurred in patient cohorts and either our sc-eVIP scores or cellular growth assays (Fig. 4a-d, TP53: Spearman correlation = 0.73, P=3.1*10^-15^ with growth in Nutlin-3, TP53-null, Spearman correlation = 0.68, P=8.1*10^-13^ with sc-eVIP scores; KRAS: Spearman correlation= 0.63, p=7.7*10^-12^ with GILA assay and Spearman correlation= 0.74, P=3.3*10^-17^ with sc-eVIP scores).

For TP53, previous work has shown that integrating functional assay results with mutational signatures accurately predicted the frequency of somatic variation across patient cohorts (Giacomelli et al., 2018). Following on this, we partitioned the variants into training and test sets by position (such that variants at the same position will not be shared between the training and test sets) 10 times, and quantified the extent to which sc-eVIP impact scores or functional assays are predictive of TP53 variant frequency, when combined with mutational signatures in a generalized linear model. All models trained performed better than random (t-test comparing each group with shuffled p. adj. <0.05), with similar performance for models using sc-eVIP scores as features, as compared to those using functional assays (Fig. 4e,f, Extended Data Figure 9a-c). As expected, models trained on the restricted set of 99 variants profiled in this work had higher variance than those trained on a larger training set, due to the model’s need for sufficient dynamic range of effect in the positions assigned to the training set.

For KRAS, models using either sc-eVIP scores or GILA scores to predict variant frequency performed significantly better than a shuffled model (t-test comparing each group with shuffled p. adj. <3.92*10^-4^) and had similar performance to each other, with the exception of models trained on a larger set of variants than present in our dataset, which had a significantly better performance than models using sc-eVIP features (p. adj. 7.51*10^-3^, Fig, 4g,h, Extended Data Figure 9d-f). Overall, variants with the largest sc-eVIP impact score in our experiment also tended to be the most commonly mutated in cancers (Fig. 4j). Other gain of function variants (*e.g.*, Q22K, A59G,E,T T58I, V14L, D119G) with somewhat lower frequencies, also had lower sc-eVIP scores within the gain-of-function cluster, as well as lower GILA scores. However, mutation frequency did not resolve well impactful variants that occur at a lower frequency (<20 observations), highlighting the distinct value of functional profiles. For example, variants at known gain-of-function hotspot positions 12, 13 and 61 that occur more rarely due to the requirement for multiple base changes (*e.g.*, G12I, G12Y, G13E, Q61A, Q61P) are still detected as impactful with our assay. Additionally, ultra rare variants requiring single base changes (such as V14L and D119G) were also found to be gain-of-function, albeit at the low end of the gain-of-function continuum. Finally, several variants that arise from single base mutation (A146P, G13R) and are comparable in frequency to the neutral group (Fig. 4j, left), score as gain-of-function in our assay (Fig. 4j, right). Together, these observations highlight interesting disconnects between mutation prevalence in patients and mutation function, highlighting the importance of generating functional data as a complementary approach to cancer genome sequencing.

### Coding variant Perturb-Seq as a scalable variant phenotyping platform

Finally, we performed power analysis by subsampling to evaluate the scalability of our approach for phenotyping thousands or more of cancer variants (Fig. 5). The impact of variants with large effect sizes could be detected with as few as 20-100 cells per variant, whereas smaller effect sizes mostly required 100-300 cells per variant (Fig. 5a-c). Thus, with ∼8.3 million single cells, one could conceivably study all ∼270,000 possible variants in each 1kb cancer gene in the Foundation Medicine Panel ([CSL STYLE ERROR: reference with no printed form.]), and with ∼71 million cells, one could create a draft cancer variant functional atlas of ∼2.3 million possible variants in the ∼200 actionable cancer genes with cDNA sizes under 3kb (Fig. 5d,e). These calculations account for a 65% rate of variant barcode detection (as observed in our study), 20 cells per variant (sufficient for detecting the largest effect sizes per variant), and would amount to 400 cells per amino acid position, allowing us to detect lower impact variants by transferring information across similar variants when modeling variant impact. As both large-scale single cell profiling and DNA synthesis technologies are rapidly increasing in scale, at lower costs (Datlinger et al., 2019; Ma et al., 2020; Sidore et al., 2019), such comprehensive experiments are now within reach.

## Discussion

Our study shows the feasibility of a scalable, general platform for variant impact phenotyping in which pooled variants across different genes can be evaluated simultaneously for function. The single unified readout of single cell gene expression does not only provide an efficient experimental approach, but is also a rich and interpretable molecular phenotype. For example, the same readout allowed us to show that the cell cycle is the main underlying signal for differences between TP53 variants, but not for KRAS variants. In KRAS, the high-dimensional, continuous, and high-resolution profiles identified a continuum from neutral to highly impactful variants and categorized this continuum into variant classes, each distributing differently along a broad phenotypic space, and associated with specific gene expression programs which shift gradually. Such quantitative shifts of the cellular landscape at the single cell level are consistent with studies on the effects of variants affecting cellular compositions (Brodin et al., 2015; Dubovik et al., 2018; Li et al., 2018).

The single cell nature of our approach allowed us to move beyond average gene expression profiles of variants to study their effect on distributions of cells. Specifically, in the cellular context explored in this work, it revealed that variants do not simply fall in discrete categories of loss-of-function or gain-of-function but rather show quantitative shifts in cell compositions. The observed re-distribution of cells across the phenotypic landscape, enriching and depleting specific cellular states for each variant, provides hypotheses for explaining the quantitative behavior of such variants in orthogonal assays. Moreover, the learned gene programs across single cells provide a step towards decoupling potentially multiple functionalities of variants for a gene, as is the case for KRAS where we observe a gain-of-function continuum as well as additional separate gene programs separating variant classes that may capture other aspects of KRAS biology to be further characterized.

While expression-based variant phenotyping can help predict mutation frequencies, some mutations present in lower frequencies can nonetheless be highly impactful in our assays, showing the power and complementarity of functional, expression-based phenotyping. In particular, certain variants in KRAS had stronger phenotypic effects than we would have predicted based on recurrence in patient samples, while others had weaker effects than we would have predicted. Such discrepancies can be explained, at least partially, by the mutability of the underlying nucleic acid residues, consistent with prior observations with TP53 (Giacomelli et al., 2018). Future work will determine if models can be trained to better incorporate mutational signatures to predict the extent of functional impact without experimental assays.

The observation that a cancer variant induces a gene expression change relative to the WT allele (defined as an impactful variant in this work) is highly suggestive of its biological function, but is not a definitive assessment of the induction of a cancer phenotype, such as tumorigenesis or drug sensitivity in models or patients. It is possible for impactful variants, as defined using gene expression profiling to not be consequential for human tumors. Nevertheless, our analysis showed high concordance between expression-based phenotyping and dedicated, highly optimized gene-specific functional assays. This suggests that an expression-based cancer variant impact atlas should be a useful first draft for annotating allele function. Comparisons to more physiological assays as they are developed would help properly calibrate false positive and false negative rates, and help interpret expression patterns in terms of their physiological relevance. With recent advances in pooled optical screens (Feldman et al., 2019), matched reference maps with both genomics and cell biology readouts should help facilitate this interpretation.

Our experimental approach can be expanded and improved in several ways. First, we focused on one cellular context, A549 lung cancer cells, but our previous work (Berger et al., 2016; Kim et al., 2016), showed that additional cellular contexts can add sensitivity regarding the mechanism of loss-of-function (*e.g.*, a WT endogenous allele is required for distinguishing dominant negative from LOF effects). The pooled nature of Perturb-Seq should allow us to readily extend this work to many cell lines, including existing pools of cell lines such as PRISM (Kinker et al., 2019; McFarland et al., 2020). Second, we used exogenously expressed cDNA constructs, where viral packaging limits the interrogation of longer (>3.5kb) genes, and where some variants may be expressed at non-physiologic levels. New approaches for base editing (Gaudelli et al., 2017; Komor et al., 2016) and prime editing (Anzalone et al., 2019) should help overcome this bottleneck. Third, we were limited by variant-level barcoding at the 3’ end of each construct for scRNA-seq detection. Advances in long-read scRNA-seq methods (Lebrigand et al., 2020; Volden and Vollmers, 2020) should allow direct sequencing readout of individual RNA variants. Finally, our scale is impacted by the costs of scRNA-seq. Advances in massively parallel scRNA-Seq, such as scifi (Datlinger et al., 2019) and combinatorial barcoding (Cao et al., 2017; Ma et al., 2020; Rosenberg et al., 2018) have substantially reduced such costs and can be efficiently applied to cell lines. Efficiency can be further enhanced by using “compressed” designs (Cleary et al., 2017), such as introducing multiple distant mutations in one construct, multiple constructs in one cell, or multiple cells profiled together.

Our proof-of-concept demonstrated the ability of sc-eVIP to read out variant impact across multiple distinct genes. The generalizability of this approach across genes will depend on whether engineered variants induce cell state changes that can be recorded at the transcriptomic level. We expect the approach to be particularly useful for variants in genes that affect many gene programs and elicit cell state changes, as for example those that result in signaling cascades or cell cycle changes. On the other hand, it may be more challenging to use gene expression based variant impact phenotyping for genes that result in transient effects or that have effect sizes comparable to the noise levels in these data. While the current work and previous research (Berger et al., 2016; Kim et al., 2016) have observed sc-eVIP to be versatile across a variety of cancer genes, future work will determine the extent to which sc-eVIP can be applied across all genes.

Overall, coding variants Perturb-Seq combined with the associated sc-eVIP analytical framework represents a versatile and powerful approach for assessing the phenotypic impact of coding variants. In contrast to existing methods, it provides a high-content and highly interpretable readout about the functional impact of overexpressed variants and does not require the development of bespoke phenotypic assays for each gene of interest. At its current scale, it can be immediately deployed to assess medium-sized libraries of hundreds of disease-related variants, important for both basic biological understanding and therapeutic applications. With improvements, it should be amenable to the assessment of tens of thousands of variants and deep mutational scans of entire disease-related genes in cancer and beyond, including coding variants in common diseases, and to construct general predictive models from variants to functions. Such an atlas of variant impact across all key cancer genes and contexts would be a foundational resource for translational cancer research.

## METHODS

### EXPERIMENTAL APPROACHES

#### Variant pool construction for TP53 and KRAS variant pools

Clinically observed variants of KRAS and TP53 were downloaded from cBioportal on October 20^th^, 2017 from the TCGA (Bailey et al., 2018), MSKCC-IMPACT (Zehir et al., 2017), and GENIE (AACR Project GENIE Consortium, 2017) pan-cancer datasets and merged into a single list. We selected the 75 most frequently observed missense alterations for each gene from this list. The goal was diversity of amino acid variants; for amino acid changes that exhibited multiple possible nucleotide alterations, we selected a single variant (with priority given to single nucleotide variants). 25 negative control variants were also selected, comprising 10 missense and 15 synonymous variants from ExAC that were not observed in the aforementioned cancer sequencing studies. We obtained wild-type TP53 (NM_000546.5) and KRAS (NM_004985.4) sequences from GenBank (Benson et al., 2013).

Variants were synthesized by Twist Bioscience, appended with unique 10 bp barcodes generated by the ‘DNAbarcodes’ R package (Buschmann and Bystrykh, 2013) using a Hamming distance of 5 between barcodes, and cloned into a modified Perturb-seq (Dixit et al., 2016) vector.

#### Perturb-seq: transduction, selection, and scRNA-Seq

10^6^ A549 cells (ATCC CCL-185) were transduced with either KRAS or TP53 lentiviral pools at an MOI of 0.1 to attain a single integration event per cell, and selected with 2 µg/mL puromycin for 2 days to obtain a final library representation of ∼1,000 cells per variant for both KRAS and TP53. Cells were allowed to recover for 2 days after selection and then loaded onto a 10X Chromium chip using the 10X Chromium Single Cell 3’ v2 kit (10X Genomics #120237). We loaded 7,000 cells per channel across 32 channels for each cDNA library to obtain a total of 224,000 cells per library (448,000 total). Paired-end libraries were sequenced over 32 lanes on an Illumina Hiseq 2500 per sequencing parameters recommended by 10X Genomics: cell barcode read length 26 bp, index read length 8 bp and transcript read length 98 bp. No dial-out PCR was done, in contrast to typical Perturb-seq (Adamson et al., 2016; Dixit et al., 2016; Jaitin et al., 2016) (Variant assignment is described in the section “Assigning variants to cells”.)

## COMPUTATIONAL ANALYSIS

### Single-cell RNA-seq data pre-processing

Sequencing reads were demultiplexed and aligned using Cellranger 2.1.1 (Zheng et al., 2017), mapping to the human transcriptome version GRCh38-1.2.0, and resulting in a matrix of Unique Molecular Identifier (UMI) counts for each gene in each cell.

We then further processed this matrix using scanpy (Wolf et al., 2018). To filter out low-quality cells and keep the most informative genes, we removed cells with <200 genes/cell, and then removed genes present in less than 3 of the remaining cells. We then further filtered out cells with fewer than 7,000 UMIs/cell and those with a percent mitochondrial UMIs/cell >20%. Finally, we down-sampled cells with >50,000 UMIs to 50,000 UMIs, to avoid cells with unusually high sequencing depth.

After filtering and the above down-sampling, to account for any differences in sequencing depth between the 64 10x channel batches, we further down-sampled the UMIs per cell such that all batches would have a median of 20,000 UMIs/cell. For this, we computed the median number of UMIs/cell in each channel, and then down-sampled the UMIs of each cell by a factor defined as the 20,000 desired UMIs/cell divided by the median number of UMIs/cell in the batch of the cell. Specifically, given a cell and this down-sampling factor, we went through each gene and obtained the adjusted number of UMIs by sampling from a binomial distribution with p=down-sampling factor and N=number of UMIs observed for this gene in the cell. This down-sampling procedure adjusted the distributions of UMIs/cell in each batch to have more similar medians. Batches with a median UMI/cell less than 20,000 did not go through this procedure. Note that in practice, similar results are obtained even without this down-sampling.

We normalized the expression UMIs per cell to sum to 10,000 in each cell, and then transformed the normalized values to log(normalized expression+1) to obtain a raw expression matrix. We selected variable genes by identifying the genes for which the variance (scaled to a z-score relative to other genes in similar expression bins) exceeded 0.5. We also filtered the variable genes to have raw expression levels between 0.0125 and 4 (Zheng et al., 2017).

We regressed out batch (as a discrete {0,1} variable), the number of UMIs/cell, the percent of mitochondrial reads, and the normalized expression of the variant barcode, and converted the resulting residuals to z-scores for each gene across cells. The analyses in this work use these z-scores, unless otherwise noted.

We performed PCA for dimensionality reduction, keeping the first 50 principal components. We represented the cells in a low dimension using UMAP, using the default values of 15 nearest neighbors per cell and the default minimum distance between embedded points of 0.5. We used the resulting nearest neighbor graph to cluster cells using Louvain clustering (Blondel et al., 2008; Levine et al., 2015).

We then subsampled the cells, to obtain 1000 for each variant, and this was used in downstream analyses.

### Investigating potential doublets

We identified Louvain clusters of cells with higher counts than the rest, and checked if there was an enrichment of cells with multiple assigned variants, or a depletion of unassigned cells, as both of these would indicate the presence of doublets (Extended Data Fig. 1g and 5g). For TP53, this flagged clusters 10 (high counts, significantly depleted in unassigned cells and significantly enriched in cells with multiple variants) and 11 (high counts, significantly depleted in unassigned cells, significantly enriched in cells with multiple variants), comprising 5.7% of all cells.

For KRAS, this flagged clusters 10 (high counts, significantly enriched in cells with multiple variants), 11 (high counts, significantly depleted in unassigned cells and significantly enriched in cells with multiple variants) and 12 (high counts, significantly depleted in unassigned cells and significantly enriched in cells with multiple variants), comprising 8.5% of cells. While we retained the cells in these clusters in the results we presented, removing them did not significantly change the results.

### Assigning variants to cells

To identify the variant(s) that were overexpressed in each cell, we identified the subset of reads that mapped to the distinct 10 bp barcodes annotating each variant. To this end, we created a reference for aligning reads consisting of the full sequence of the plasmid used in this study. We aligned all scRNA-seq reads to this reference, using Bowtie (version 1.2.2 (Langmead et al., 2009)), and conservatively only kept alignments with up to 2 mismatches. We kept only valid alignments, defined as aligning to the negative strand of the reference (and positive strand for reads that align to the puromycin resistance gene on the construct). We filtered out UMIs with a transcript-per-transcript (TPT) value < 1 (Dixit, 2016), to remove chimeric reads containing variant barcodes. TPT quantifies for each read in a combination of cell barcode and UMI the fraction of reads that are identical, in order to flag cases where for a given cell-barcode-UMI combination, there is a small fraction of reads that are chimeric and do not map to the same variant barcode as the majority of reads. We considered reads having the same cell barcode and UMI and mapping to the same variant barcode as identical, even if they mapped to a different position of the plasmid, reasoning that they may still come from the same original molecule. We assigned cells to variants as long as there was at least 1 read supporting the variant. Because we use even single reads to assign variants, it is possible that we are overestimating the number of cells with more than 1 variant (and underestimating the number of cells that contain only a single variant).

### Sc-eVIP impact scores

To quantify the degree to which expression profiles of cells overexpressing a variant deviate from those of cells overexpressing the wildtype version, we used a Hotelling’s T^2^ statistic, a multivariate generalization of a *t*-test (Hotelling, 1931). We compared the profiles of cells overexpressing WT *vs*. mutant proteins in principal component space, considering the top 20 PCs, with the T^2^ value reported as the test statistic. Higher scores indicate higher impact.

We then derived an empirical null distribution of the scores and identified a threshold corresponding to a False Discovery Rate of 1%, as previously described (Noble, 2009). When we only simply permuted the assignments of the variants to the cells, control synonymous variants scored as impactful (Extended Data Fig. 1h, 5h), because of variation between synonymous variants. We thus derived our empirical null distribution by comparing pairs of control synonymous variants in our dataset, and computing the FDR at a given score threshold as: (percent scores higher than the threshold from the empirical distribution)/(percent scores higher than the threshold from the empirical distribution + percent scores of non-synonymous variants higher than the threshold). Finally, we identified the highest score threshold associated with our desired FDR of 1%.

### Clustering variants by mean expression profiles

To cluster variants into discrete classes, we computed the Spearman correlation coefficient between the average gene expression profiles of each pair of variants and used the correlation matrix as an input for clustering using an *L*1 distance and complete linkage. We ordered the leaves of the resulting dendrogram by increasing effect sizes, as computed with the Hotelling T^2^ test. To obtain discrete cluster assignments, we cut the hierarchy based on visual inspection, obtaining 3 clusters for TP53 and 5 for KRAS.

### Computing gene programs

To identify genes whose expression is impacted by variants, we clustered the genes by variants matrix of average expression profiles across variants using Louvain clustering (using 5 nearest neighbors). This yielded sets of genes with coherent average expression profiles across variants. To identify representative genes for each such gene program, we asked which genes are most correlated with average program scores across variants. For this, we averaged the expression of genes in each program across in each cell, then averaged scores across cells of a given variant and then finally computed the correlation between the average gene expression program and the average expression of a gene across variants.

### Scoring of cell cycle phases in single cells

To identify the cell cycle phase of each cell, we followed our previously described approach (Macosko et al., 2015). Briefly, we retrieved the representative genes for each of the cell cycle phases G1/S, S, G2/M, M, M/G1. We then restricted the sets of genes in each cell cycle phase to those that were correlated with the overall score (Spearman correlation coefficient > 0.3). We scored each cell cycle phase, and standardized the scores to z-scores within each cell cycle phase. We then assigned cells to the phase that had the highest score. If a cell had low scores for all phases (z-scores < 0), it was classified as non-cycling.

### Signature analyses

To compare our results with previously reported signatures for TP53, we retrieved two signatures from (Fischer, 2017; Jeay et al., 2015), consisting of two lists of genes. For each signature, we scored each cell by computing the average score across the genes in the signature, relative to a set of 50 control genes, chosen at random in a manner stratified by expression levels to match those of the genes in the signature. We then averaged signature scores across all cells for each variant and subtracted the average score in unassigned cells to obtain the results presented in Fig. 1g.

### Comparison to dedicated cellular variant phenotyping assays

We retrieved TP53 cellular variant phenotyping assay data from (Giacomelli et al., 2018). For each of three conditions (Nutlin-3 in TP53-WT cells, Nutlin-3 in TP53-null cells and etoposide in TP53-null cells) in A549 cells, the values retrieved represent the fitness change induced by overexpressing the variant after 12 days of treatment, reported as z-scores of log2-fold changes between the number of cells with the variant at the end of treatment and those at the beginning.

We retrieved KRAS growth in low attachment (GILA) measurements from (Ly, 2018). The measurements were done on HA1E cells at 2 timepoints, 7 and 14 days, and were reported as z-scores across all variants tested. The two timepoints were highly concordant (we report Day 7 results in the main text, and show both in Extended Data Fig. 7a-b).

### Models to predict the identity of variants based on gene expression

We trained a multi-class logistic regression classifier to distinguish each variant cluster for each of TP53 and KRAS, using the method *sklearn.linear_model.LogisticRegression* from the package *sklearn* in python, with loss set to ‘multinomial’. We balanced the number of examples per class by subsampling each class of variants to the class with the fewest cells (6000 cells per class for TP53 and 3000 cells per class for KRAS). We partitioned the cells or the variants depending on the task into 50% in the training set and 50% test sets. We computed AUPRC using the R package PRROC (Grau et al., 2015).

### Studying compositional changes induced by variants

Given an assignment of cells to sets, either cell cycle phases or clusters, we tested for each variant whether the distribution of cells from this variant across groups differs from that of cells overexpressing the wildtype allele, quantifying significance using a chi-square test.

### Predicting mutation frequencies in cancer cohorts

To predict mutation frequency of a variant across patient cohorts, we followed a previously described procedure (Giacomelli et al., 2018). Briefly, we retrieved mutational signature scores and cellular variant phenotyping assay data (Giacomelli et al., 2018) and then fit a generalized linear model to predict the counts of each variant in cancer cohorts as measured by IARC counts for TP53. We used the package statsmodels in python, with the command *statsmodels.GLM*, with *family=sm.families.Poisson()*). The features used were combinations of: (**1**) impact scores, (**2**) mutational signatures and (**3**) cellular variant impact phenotyping assays from (Giacomelli et al., 2018). Given the small number of examples for training, we used only the mutational signatures 1, 2, 4, 5, 6, 7, 13, 24 as these were previously deemed most informative (Giacomelli et al., 2018).

We partitioned the data into training and test sets for TP53 by amino acid position, to avoid train-test contamination through different variants at the same position having similar effects. For each type of model, we trained 10 cross-validated models, and report the median performance in the main text, and show the distribution of performance scores (Fig. 4f,h, Extended Data Fig. 9a-c for TP53, and Extended Data Fig. 9d-f for KRAS). For sc-eVIP impact scores, the training and test sets contained only the variants profiled in this study. For the comparisons with functional assays and mutational signatures, we also trained on the full datasets from (Giacomelli et al., 2018). As a control, we also consider the performance obtained when shuffling the order of the true counts. We focused on the subset of variants annotated with mutational signatures, which resulted in the control synonymous variants being excluded from these analyses.

We used a similar approach for the variants in KRAS. We excluded from the training variants with 0 observed occurrences in cancer cohorts.

As performance metrics, we report Spearman and Pearson correlation coefficients between the ground truth and the predicted values, as well as R^2^ (coefficient of determination) as computed with the *sklearn* package in python, with the function *sklearn.metrics.r2_score* (note that this score can be negative, if a model predicts worse than a model predicting the average of observed values).

### Power analyses

To determine the impact of the number of cells profiled per variant and the variant’s strength on the ability to detect variant impact, we performed a cell subsampling experiment, and quantified performance as the fraction of impactful variants with a given effect size (as determined from the full dataset) that are recovered at an FDR of 1%, for the number of cells allotted. We performed 10 independent subsampling iterations, and report the average performance for a given impact score and number of cells per variant.

To ensure comparable results between TP53 and KRAS, for which we had a limited number of cells overexpressing wildtype alleles, we used synonymous variants that had >=1,000 cells/variant as the “WT reference” (P359P for TP53 and K169K for KRAS), rather than the WT overexpressing cells.

### Projections for creating an atlas of cancer variant impact

We computed the number of cells and associated costs for characterizing all possible variants in a set of actionable cancer genes, which we retrieved from the Foundation Medicine Panel ([CSL STYLE ERROR: reference with no printed form.]). For each gene, we used the APPRIS database (Rodriguez et al., 2018) to select a principal isoform to serve as the basis for our calculations. We computed the number of variants for a given gene by multiplying the number of codons in its ORF by 20 amino acids. The number of required cells was defined as the number of variants multiplied by 20 cells per variant (a tradeoff to directly detect alleles with the strongest effect, as well as pool data from different variants in one position for alleles with smaller effect sizes). We increased the number of required cells (by 1/0.65), to account for our average detection rate of 65% cells with a single variant.

### Reported p-values

P-values reported throughout the paper are adjusted p-values, with the procedure by Benjamini-Hochberg, computed using the python package *statsmodels*, with the function *multipletests* and with the parameter method set to “fdr_bh”.

### Variant-by-variant analyses

For a comprehensive view of all analyses, displayed for each specific variant, refer to Extended Data Figures 4 and 8.

### Code availability

All analyses can be recapitulated with Jupyter notebooks at https://github.com/klarman-cell-observatory/sc_eVIP, and using the Perturb-seq library at https://github.com/klarman-cell-observatory/perturbseq.

## AUTHOR CONTRIBUTIONS

Conceptualization, J.S.B, A.H.B, J.T.N, and A.R.; Software, O.U.; Formal Analysis, O.U., and L.J-A.; Investigation, E.S., P.I.T., L.N., D.D., C.D., J.B., M.M.M., C.F., S.H.L., O.R-R. and J.T.N.; Writing – Original Draft, O.U., J.T.N., A.R., and J.S.B.; Writing - Review and Editing, O.U., J.T.N., J.S.B., A.R., W.C.H., A.O.G., A.J.A., J.B., E.S, P.I.T., and A.H.B.; Supervision, JTN, O.R-R, A.R., and J.S.B.; Funding Acquisition, J.T.N., W.C.H., O.R-R, A.R., and J.S.B.

## ACKNOWLEDGEMENTS

We thank Leslie Gaffney for help with figure preparation. We thank Alex Wu for sharing code for cell cycle analyses and Adam Rubin, Gokcen Eraslan, Brian Cleary, Elena Torlai-Triglia, Dana Silverbush, Kathryn Geiger-Schuller, Hattie Chung and Kirk Gosik for helpful discussions. We acknowledge the American Association for Cancer Research and its financial and material support in the development of the AACR Project GENIE registry, as well as members of the consortium for their commitment to data sharing. Interpretations are the responsibility of study authors. L.J.A. was a fellow of the Eric and Wendy Schmidt postdoctoral program and a CRI Irvington Fellow supported by the CRI. L.J.A. holds a Career Award at the Scientific Interface from BWF. P.T. is supported by a NIH F32 fellowship F32AI138458. Work was supported by NIH grant U01 CA176058 (J.S.B., W.C.H.), a Broad*next*10 internal award (J.S.B), a Broad Variant-to-Function (V2F) award (J.T.N.), and the Klarman Cell Observatory, HHMI, and NHGRI CEGS (all to A.R.).

## CONFLICT OF INTEREST STATEMENT

A.R. is a founder and equity holder of Celsius Therapeutics, an equity holder in Immunitas Therapeutics and until July 31, 2020 was an SAB member of Syros Pharmaceuticals, Neogene Therapeutics, Asimov and ThermoFisher Scientific. From August 1, 2020, A.R. is an employee of Genentech. W.C.H. is a consultant for ThermoFisher, Solvasta Ventures, MPM Capital, KSQ Therapeutics, iTeos, Tyra Biosciences, Frontier Medicine, Jubilant Therapeutics and Paraxel. A.O.G. is a shareholder of 10x Genomics. A.R., P.T., J.T.N, J.B. and O.U. are named co-inventors on a patent application related to sc-eVIP (US20200283843A1).

---

**Extended Data Figure 1.**
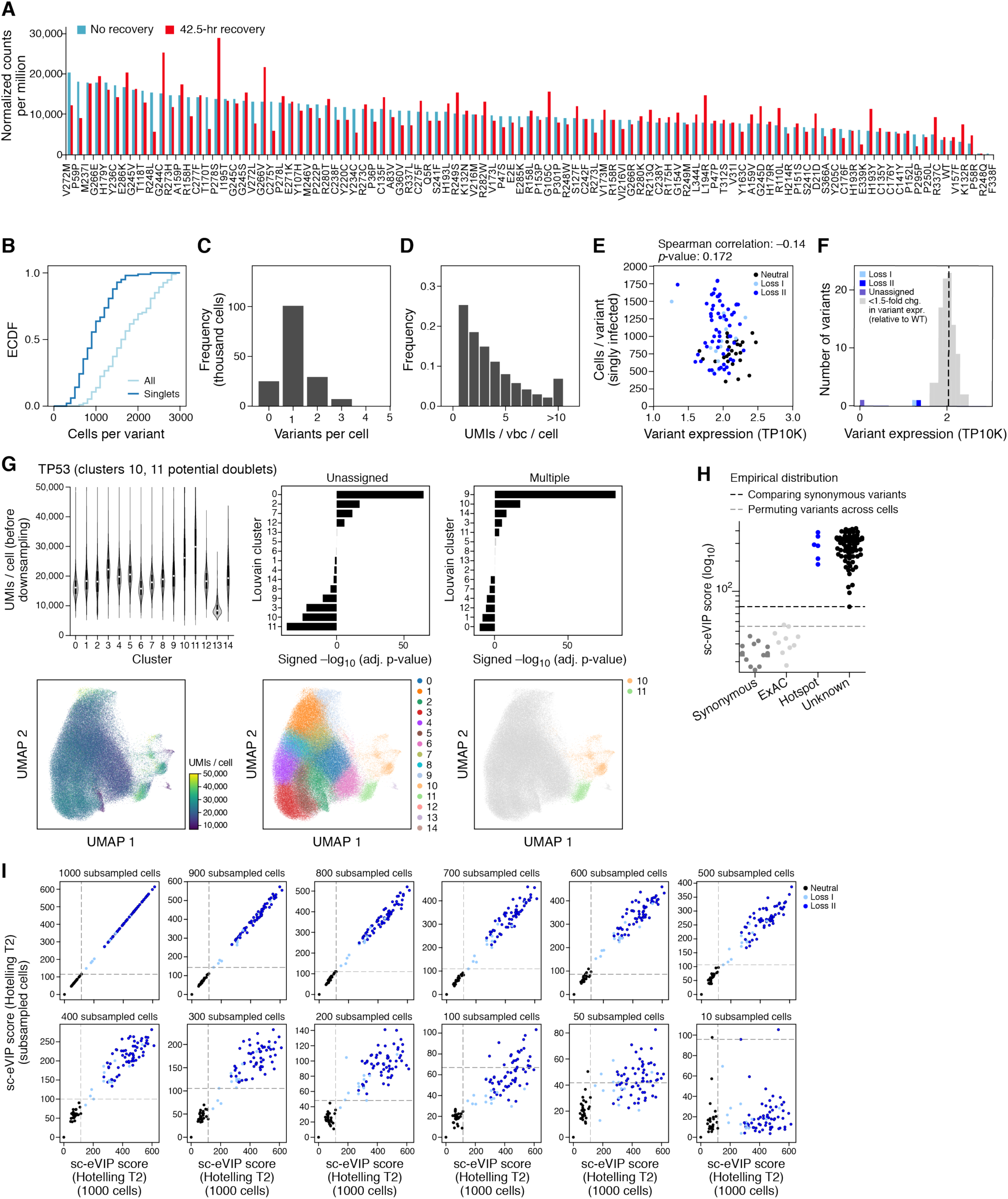
Quality control for TP53 Perturb-Seq experiment. **a.** Variant representation in the library. Number of barcode reads (*y* axis) for each tested variant (x axis), either after transduction and 2-day puromycin selection (“no recovery”), or 42.5 hours after puromycin selection (“42.5h recovery”). **b-f**. Quality control metrics. **b**. Cumulative distribution function (CDF) of number of cells (*x* axis) profiled for each variant, considering either all cells (light blue) or only cells with a single variant (dark blue). **c**. Distribution of the number of variants detected per cell. **d**. Distribution of the number of variant barcode (vbc) UMIs per cell per variant. **e**. The number of cells detected per variant (*y* axis) and the variant’s barcode expression (*x* axis, TP10K) for cells with a single variant, colored by class. **f**. Distribution of mean variant barcode expression (TP10K, *x* axis). Variants with a fold change higher than 1.5 compared to the WT barcode are colored by variant class. **g.** Potential doublets. Top left: Distribution of number of UMIs/cell (y axis) for each cell cluster as defined by Louvain clustering of cell profiles (bottom, middle). Top middle and right: enrichment and depletion (x axis, -log_10_(adj. p-value), hypergeometric test); positive sign for enrichment, and negative for depletion) for each cluster in unassigned cells (top middle) and cells with multiple variants (top right). Bottom: UMAP embedding of single cell profiles (dots) colored by number of UMIs/cell (left), cluster assignment (middle), or cell clusters that are likely doublets (clusters 10 and 11, right). **h**. Permutation tests for FDR control. Sc-eVIP score (*y* axis) for variants (dots) in each variant group (x axis). Dotted lines: FDR 1% for a permutation test shuffling the assignments of variants (gray) or estimating an empirical distribution of the sc-eVIP under the null hypothesis using only comparisons between synonymous variants (black). **i**. Impact of number of cells on sc-eVIP scores. sc-eVIP scores for each variant (dots, colored by variant class) computed using 1,000 cells/variant (*x* axis) or with varying numbers of subsampled cells (*y* axis). Dotted lines: threshold sc-eVIP score at a 1% FDR for each axis.

**Extended Data Figure 2.**
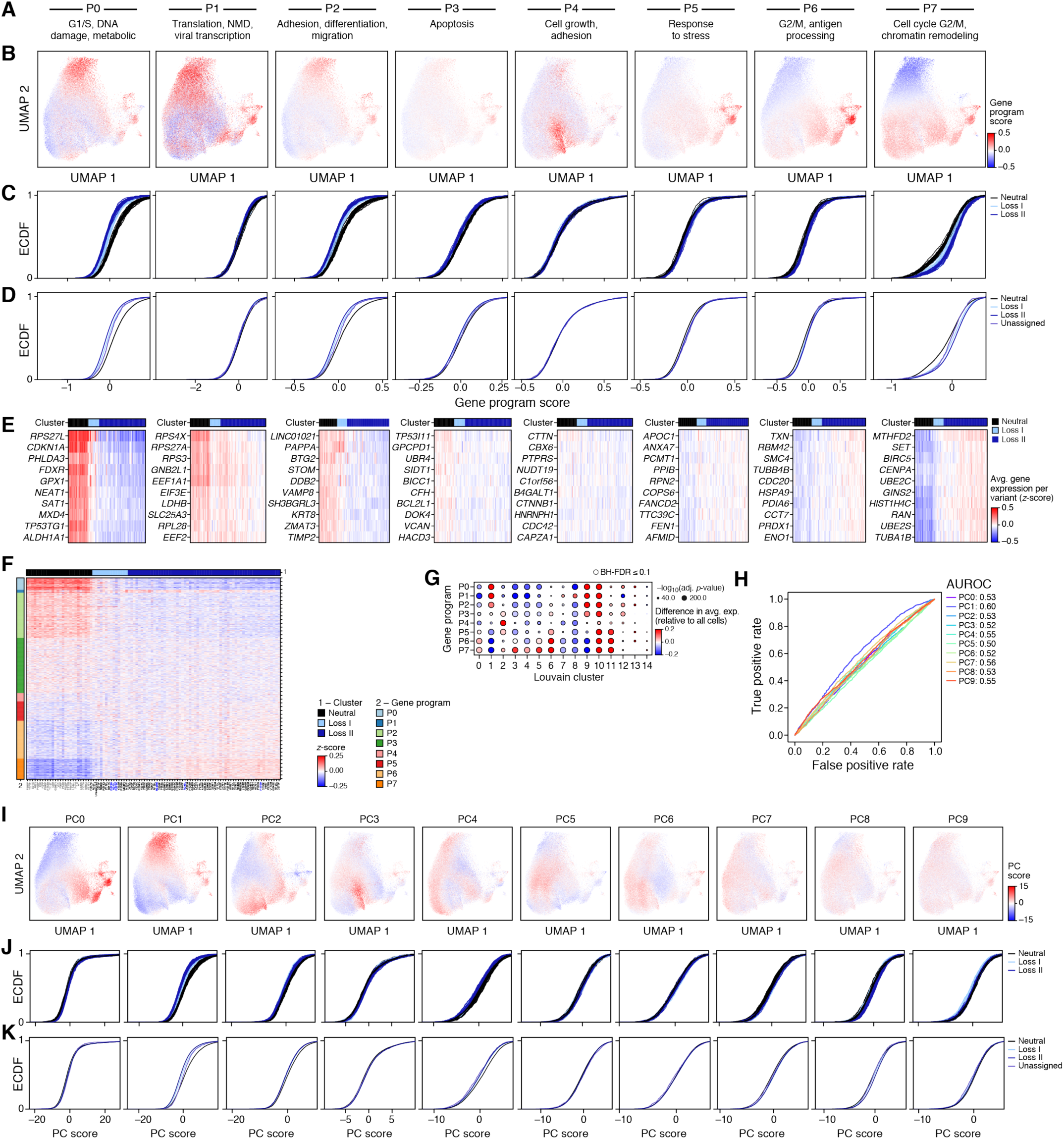
Gene programs impacted by TP53 variants. **a-e**. Gene programs impacted by variant classes. **a,b.** UMAP embedding of single cell profiles (dots), colored by program scores (color bar) and labeled by selected Gene Ontology biological processes enriched in genes from each program (top). **c,d**. CDF for program scores (*x* axis) for each variant (c) or for all variants in one class (**d**), colored by class. **e**. Average expression (z score, color bar) in cells of each variant (columns) of genes (rows) most correlated with the mean of the expression program. **f**. Mean expression (colorbar) of each gene (rows) in cells of each variant (columns). Row color bar: gene program membership; Column color bar: variant class. **g**.. Difference (dot color) in mean expression of each gene program (rows) between the cells in each cluster (columns, as in Extended Data Fig. 1g) and all other cells, and the significance of this difference (dot size, -log_10_(adj. p-value), Kolmogorov-Smirnov test, **Methods**). Colored border: BH FDR<10%. h. ROC curve of the true positive (y axis) and false positive (x axis) rate when using each PC (color) to distinguish between single cells with synonymous variants and those with variants in hotspot positions 175, 248, and 273. Color legend: Area Under the ROC curve (AUROC) for each variant. **i-k.** Principal component analysis. **i**. UMAP embedding of single cell profiles, colored by principal component (PC) scores (color bar), for each of the first 10 PCs. **j,k**. CDFs for the PC scores (*x* axis) for the cells of each variant or all variants in a class, colored by class.

**Extended Data Figure 3.**
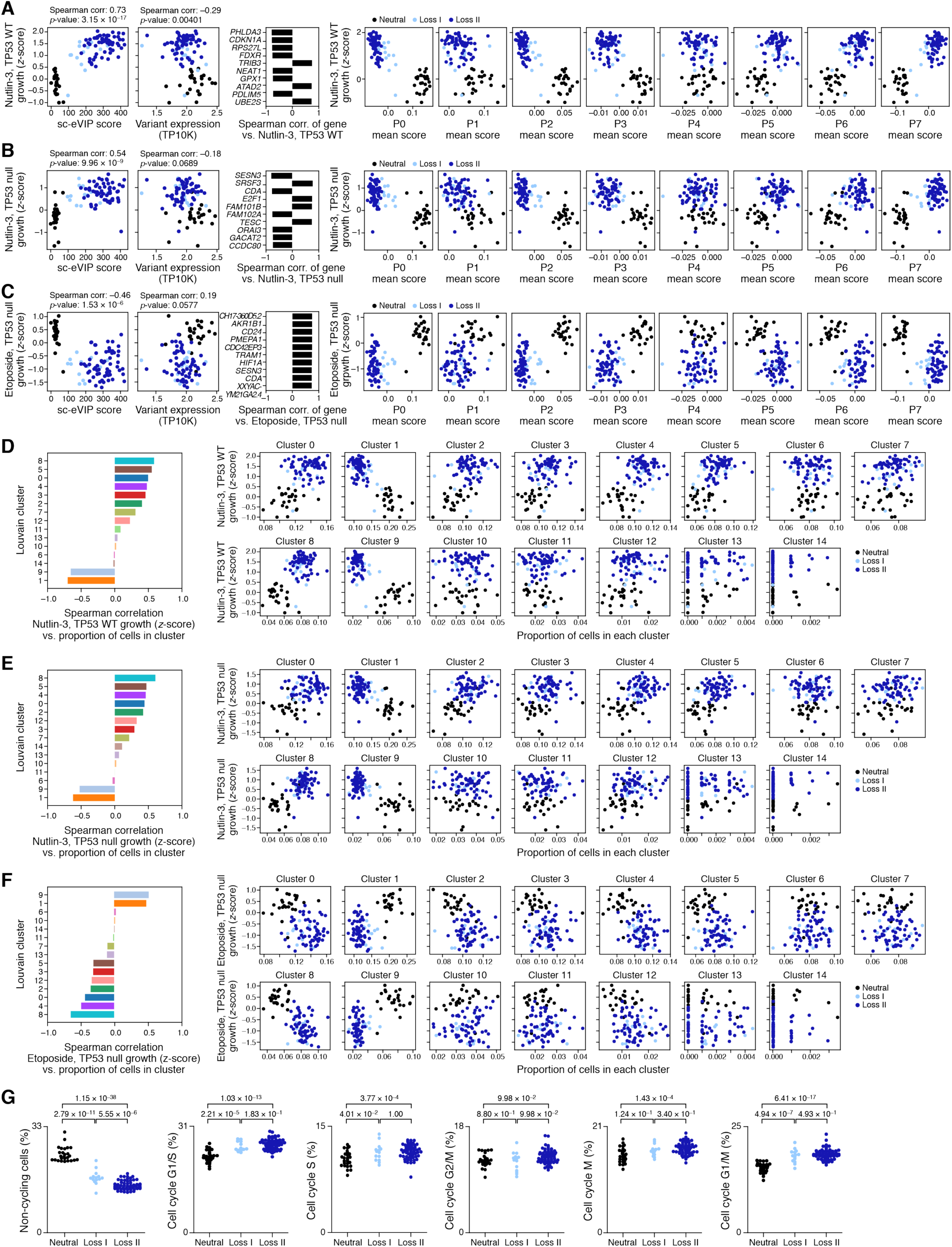
Comparison of sc-eVIP with cellular phenotyping assays and cell cycle effects for TP53 variants. **a-c**. sc-eVIP impact scores and gene programs agree with functional growth assays under Nutlin-3 treatment in a p53 wildtype background (a) or a p53 null background (**b**) and under etoposide in a p53 null background (**c**). Left: Functional assay score (*y* axis) and sc-eVIP score (*x* axis, left) or normalized variant expression (*x* axis, transcripts per 10,000 UMIs/cell (TP10K), right) for each variant (dots), colored by variant class. Middle: Correlation (*x* axis) between the functional assay score for each variant and mean gene expression across the variants for the genes (*y* axis) whose expression is most strongly correlated with the functional assays score. Right: Mean gene program (*x* axis) and functional assay (*y* axis) scores for each variant (dots), colored by variant class. **d-f**. Cell clusters correlated with functional growth assays under Nutlin-3 treatment in a p53 wildtype background (**d**) or a p53 null background (**e**) and under etoposide in a p53 null background (**f**). Left: Spearman correlation coefficient (x axis) between the proportion of cells from each variant in each cluster (as in Extended Data Fig. 1g) and the functional assay scores of the variants. Right: Proportion of cells (x axis) in cluster (label, top) and functional assay scores (*y* axis) for each variant (dots), colored by variant class. **g**. Proportion of cells in each cell cycle phase (*y* axis) among cells carrying each variant (dots) across variant classes (*x* axis). Adj. p-value: t-test.

**Extended Data Figure 4.**
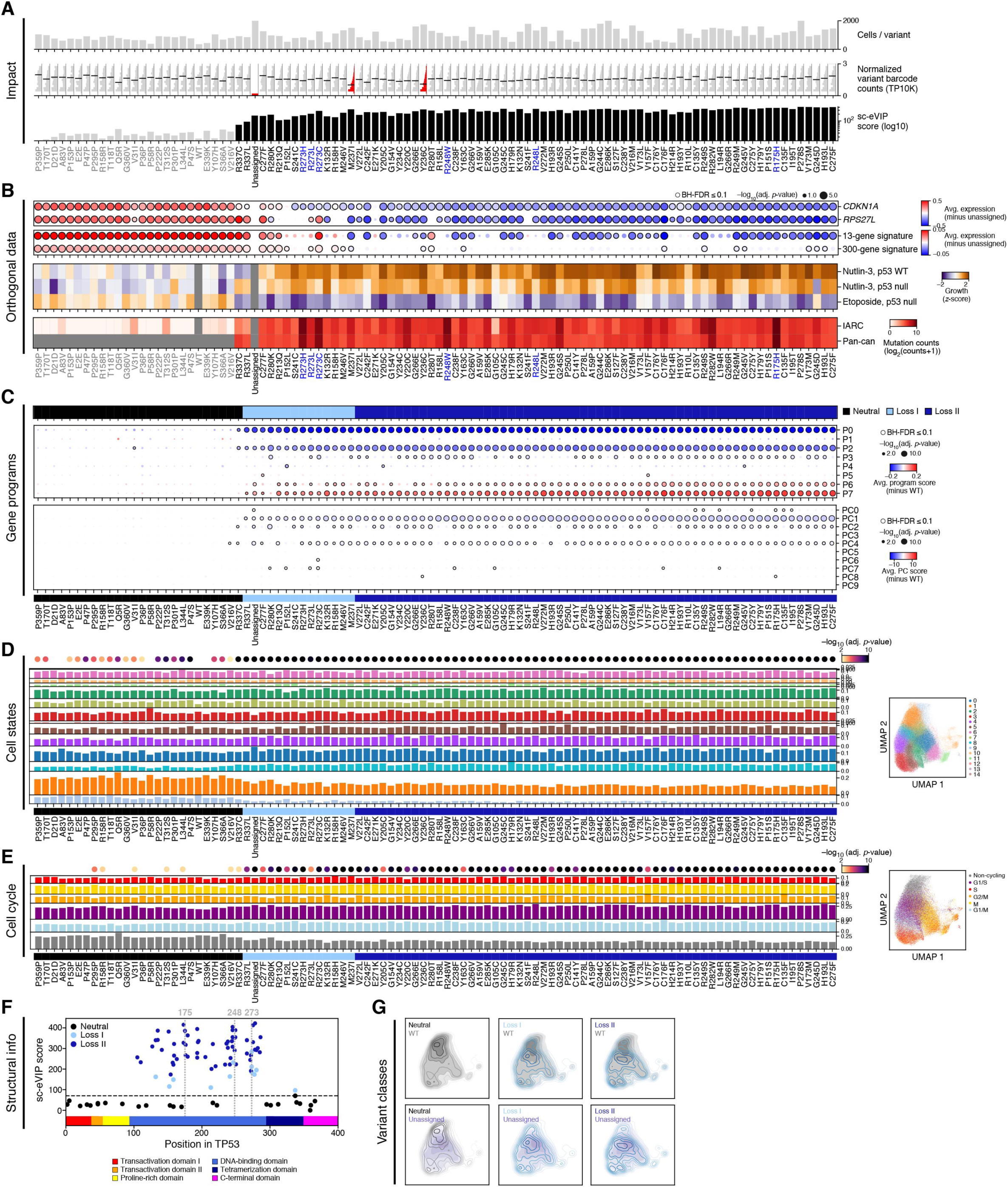
Variant-by-variant analyses for TP53 variants. **a**. Variant features. Number of cells (*y* axis, top), distribution of normalized variant barcode expression *(y* axis, middle; red: variants with a fold-change greater than 1.5) and sc-eVIP scores (*y* axis, bottom; black: significant scores) for each variant (x axis), ordered as in Fig. 1e. Grey font: controls (synonymous and ExAC), blue font: hotspot variants (positions 175, 248, 273). **b**. Agreement with other data features. Top: difference (dot color) in mean expression or signature score between a variant (columns, ordered as in Fig. 1e) and unassigned cells and the significance of this difference (dot size, -log_10_(adj. p-value), Kolmogorov-Smirnov test, **Methods**) for each of two genes canonically induced by TP53 and two TP53-associated signatures (rows). Colored border: BH FDR<10%. Middle: Growth (z-score, color bar) in three functional assays (rows) of each variant (columns). Bottom: Mutation prevalence (log_2_(counts+1) of variant occurrences) in two datasets (rows) of each variant, ordered as in Fig. 1e. **c**. Gene programs association with variants. Top: Difference (dot color) in mean program score (top) or mean PC score (bottom) between a variant (columns) and WT overexpressing cells and the significance of this difference (dot size, -log_10_(P-value), Kolmogorov-Smirnov test, **Methods**) for each gene program (top, rows, by gene clustering genes, **Methods**), or each of the top 10 PCs (bottom, rows). Colored border: BH FDR<10%. **d,e**. Relation of variants to different clusters and cell cycle phases. Left: Proportion of cells (bar height) in each cell cluster (**d,** as in Extended Data Fig. 1g) or cell cycle phase (**e**) (rows) derived for each variant (columns), annotated at the top with significance from a chi-square test comparing the cell state distribution of each variant with that of WT overexpressing cells (-log_10_(adj. p-value)). Right: UMAP embedding of single cell profiles, colored by cell clusters (**d**) or cell cycle phase (**e**). **f**. Relation of variants to p53 gene structure. sc-eVIP scores (*y* axis) of each variant (dot, colored by the variant class) and its position along the TP53 gene (*x* axis, annotated by domain). **g**. Variant induced shift in cell distributions. Density map of cell profiles in a UMAP embedding, comparing the density of cells overexpressing variants in each of 3 classes to either the WT TP53 allele (grey, top) or unassigned cells (purple, bottom).

**Extended Data Figure 5.**
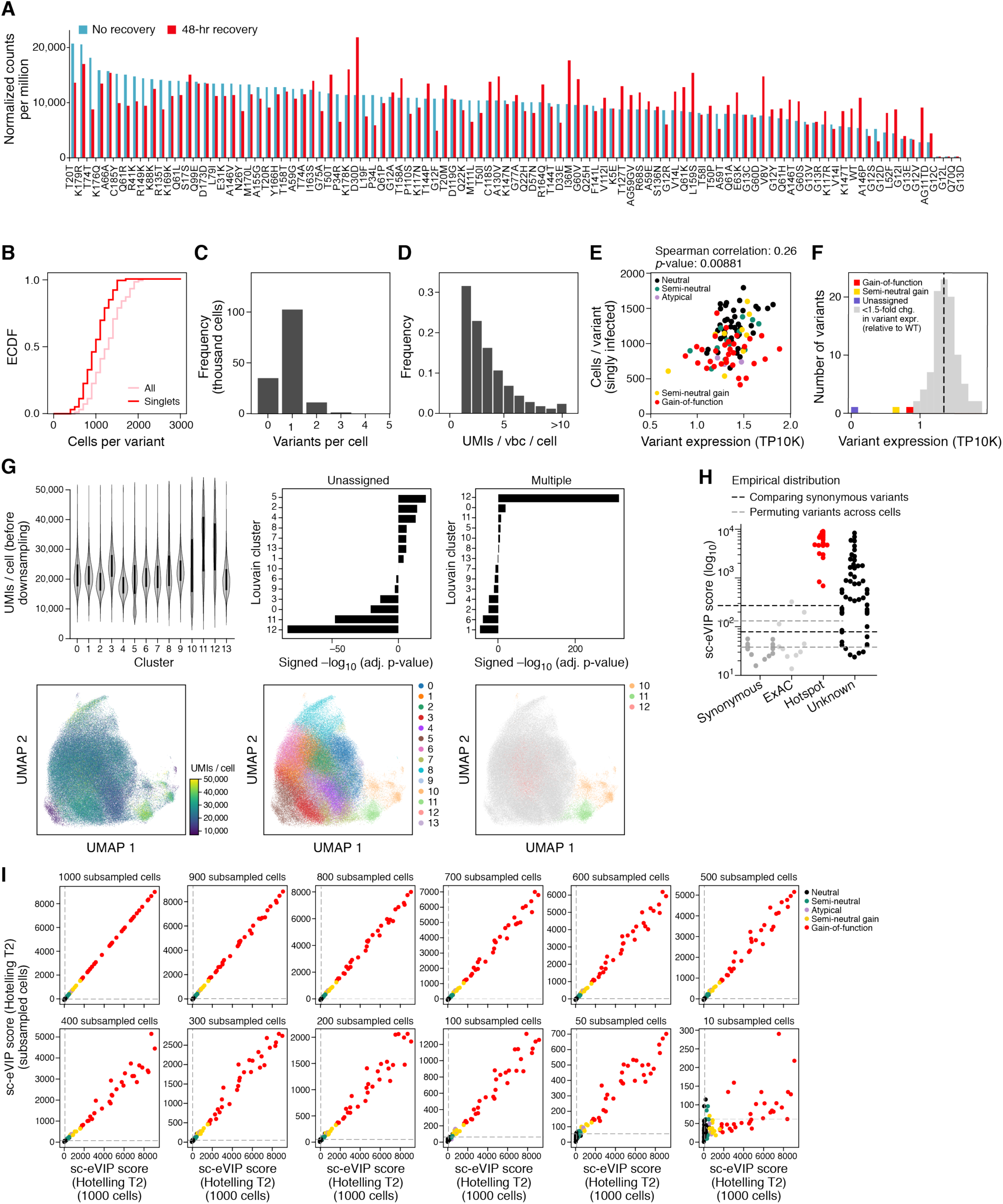
Quality control for KRAS Perturb-Seq experiment. **a.** Variant representation in the library. Number of barcode reads (y axis) for each tested variant (x axis), either after transduction and 2-day puromycin selection (“no recovery”), or 42.5 hours after puromycin selection (“42.5h recovery”). **b-f**. Quality control metrics. **b**. Cumulative distribution function (CDF) of number of cells (*x* axis) profiled for each variant, considering either all cells (pink) or only cells with a single variant (red). **c**. Distribution of the number of variants detected per cell. **d**. Distribution of the number of variant barcode (vbc) UMIs per cell per variant. **e**. The number of cells detected per variant (*y* axis) and the variant’s barcode expression (*x* axis, TP10K) for cells with a single variant, colored by class. **f**. Distribution of mean variant barcode expression (TP10K, *x* axis). Variants with a fold change higher than 1.5 compared to the WT barcode are colored by variant class. **g.** Potential doublets. Top left: Distribution of number of UMIs/cell (y axis) for each cell cluster as defined by Louvain clustering of cell profiles (bottom, middle). Top middle and right: enrichment and depletion (x axis, -log_10_(adj. p-value), hypergeometric test); positive sign for enrichment, and negative for depletion) for each cluster in unassigned cells (top middle) and cells with multiple variants (top right). Bottom: UMAP embedding of single cell profiles (dots) colored by number of UMIs/cell (left), cluster assignment (middle), or cell clusters that are likely doublets (clusters 10, 11 and 12, right). **h**. Permutation tests for FDR control. Sc-eVIP score (*y* axis) for variants (dots) in each variant group (x axis). Dotted lines: FDR 1% for a permutation test shuffling the assignments of variants (gray) or estimating an empirical distribution of the sc-eVIP under the null hypothesis using only comparisons between synonymous variants (black). **i**. Impact of number of cells on sc-eVIP scores. sc-eVIP scores for each variant (dots, colored by variant class) computed using 1,000 cells/variant (*x* axis) or with varying numbers of subsampled cells (*y* axis). Dotted lines: threshold sc-eVIP score at a 1% FDR for each axis.

**Extended Data Figure 6.**
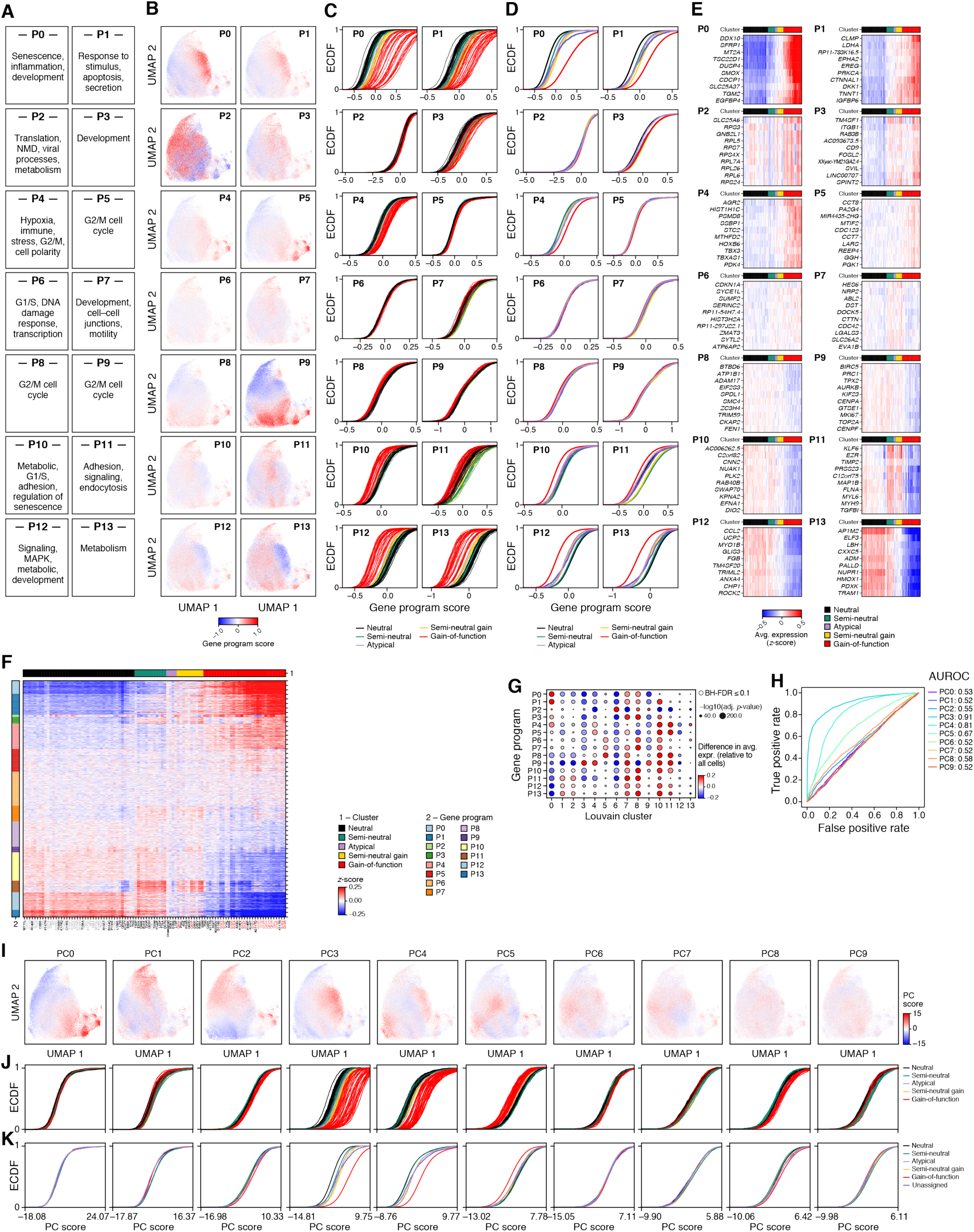
Gene programs impacted by KRAS variants. **a-e**. Gene programs impacted by variant classes. **a,b.** UMAP embedding of single cell profiles (dots), colored by program scores (color bar) and labeled by selected Gene Ontology biological processes enriched in genes from each program (top). **c,d**. CDF for program scores (*x* axis) for each variant (c) or for all variants in one class (**d**), colored by class. **e**. Average expression (z score, color bar) in cells of each variant (columns) of genes (rows) most correlated with the mean of the expression program. **f**. Mean expression (colorbar) of each gene (rows) in cells of each variant (columns). Row color bar: gene program membership; Column color bar: variant class. **g**. Difference (dot color) in mean expression of each gene program (rows) between the cells in each cluster (columns, as in Extended Data Fig. 1g) and all other cells, and the significance of this difference (dot size, -log_10_(adj. p-value), Kolmogorov-Smirnov test, **Methods**). Colored border: BH FDR<10%. h. ROC curve of the true positive (y axis) and false positive (x axis) rate when using each PC (color) to distinguish between single cells with synonymous variants and those with variants in hotspot positions 12, 13 and 61. Color legend: Area Under the ROC curve (AUROC) for each variant. **i-k.** Principal component analysis. **i**. UMAP embedding of single cell profiles, colored by principal component (PC) scores (color bar), for each of the first 10 PCs. **j,k**. CDFs for the PC scores (*x* axis) for the cells of each variant or all variants in a class, colored by class.

**Extended Data Figure 7.**
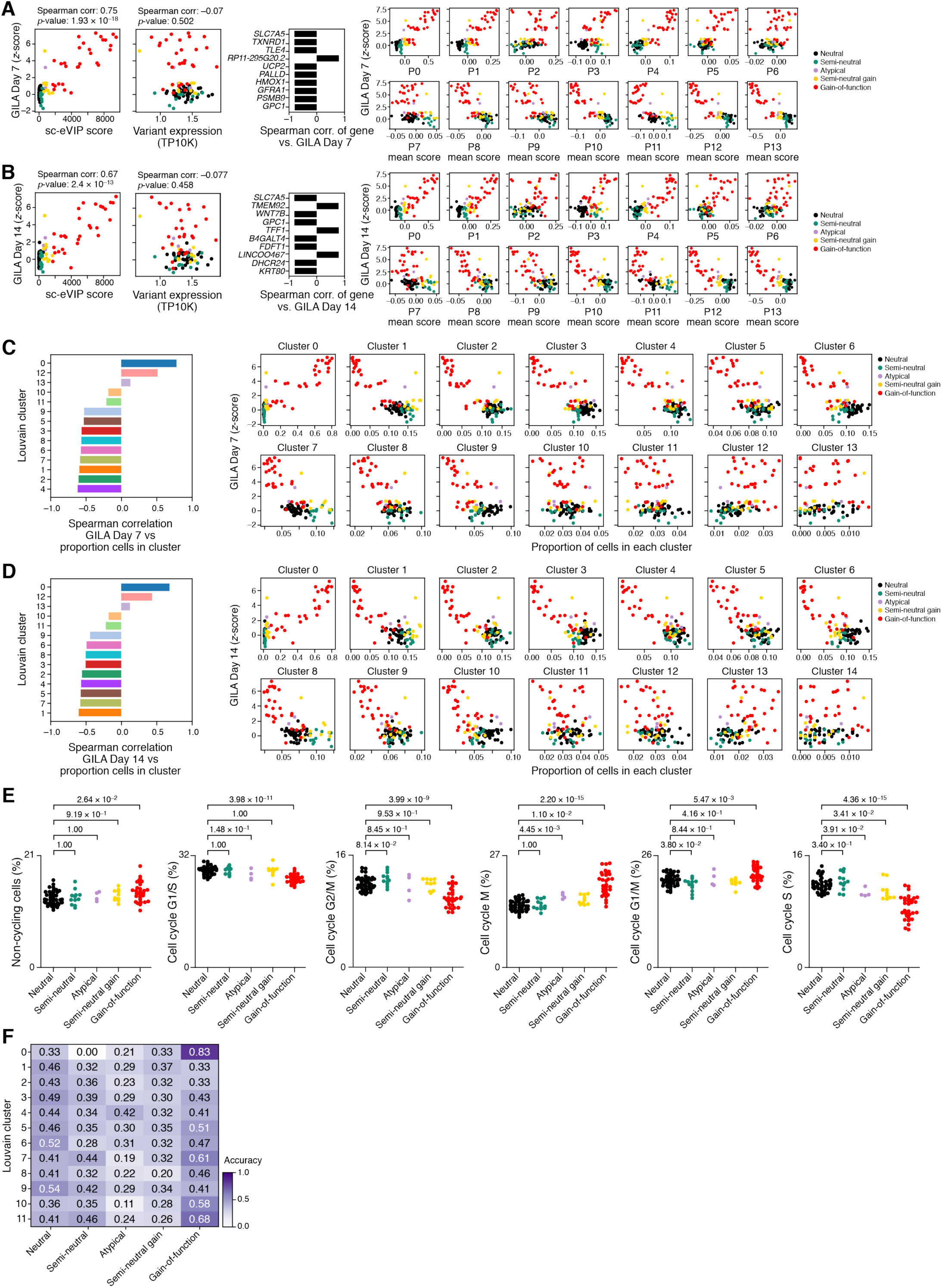
Comparison of sc-eVIP with cellular phenotyping assays and cell cycle effects for KRAS variants. **a,b**. sc-eVIP impact scores and gene programs agree with growth in low attachment at 7 days (**a**) and 14 days (**b**). Left: GILA score (*y* axis) and sc-eVIP score (*x* axis, left) or normalized variant expression (*x* axis, transcripts per 10,000 UMIs/cell (TP10K), right) for each variant (dots), colored by variant class. Middle: Correlation (*x* axis) between the GILA score for each variant and mean gene expression across the variants for the genes (*y* axis) whose expression is most strongly correlated with the GILA score. Right: Mean gene program (*x* axis) and GILA (*y* axis) scores for each variant (dots), colored by variant class. **c,d**. Cell clusters correlated with GILA at 7 days (**c**) and 14 days (**d**). Left: Spearman correlation coefficient (x axis) between the proportion of cells from each variant in each cluster (as in Extended Data Fig. 5g) and the GILA scores of the variants. Right: Proportion of cells (x axis) in cluster (label, top) and GILA scores (*y* axis) for each variant (dots), colored by variant class. **e**. Proportion of cells in each cell cycle phase (*y* axis) among cells carrying each variant (dots) across variant classes (*x* axis). Adj. p-value: t-test. **f**. Performance of a logistic regression classifier, trained to predict for each individual cell its variant class. The performance is shown as a heatmap, for each variant class (*x* axis) as a function of cell states (*y* axis), with values representing the accuracy within the cells in the respective variant class and cell state.

**Extended Data Figure 8.**
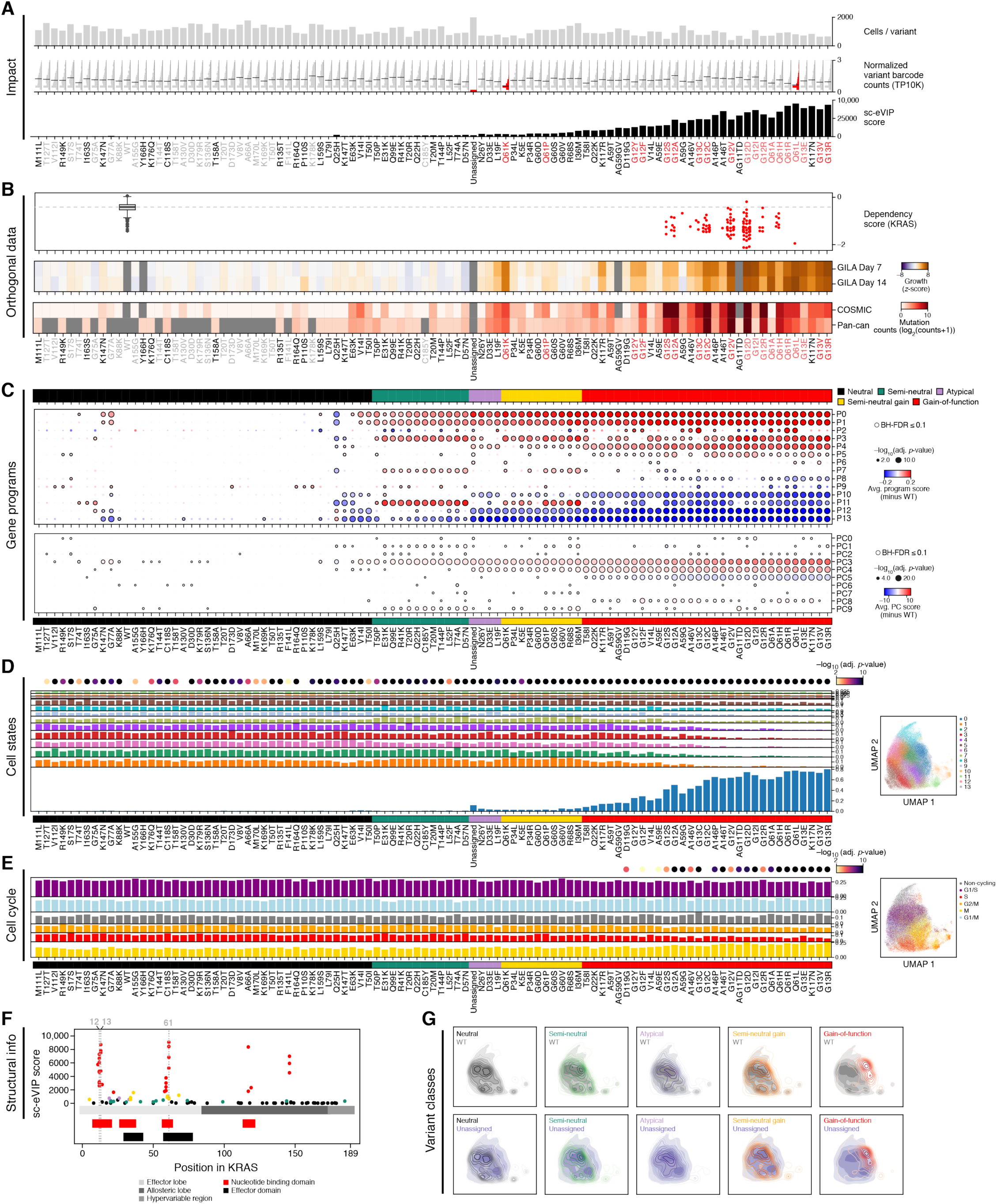
Variant-by-variant detailed representation of all analyses for KRAS variants. **a**. Variant features. Number of cells (*y* axis, top), distribution of normalized variant barcode expression *(y* axis, middle; red: variants with a fold-change greater than 1.5) and sc-eVIP scores (*y* axis, bottom; black: significant scores) for each variant (x axis), ordered as in Fig. 2d. Grey font: controls (synonymous and ExAC), red font: hotspot variants (positions 12, 13 and 61). **b**. Agreement with other data features. Top: Dependence of cell line growth on KRAS (*y* axis), for cell lines (dots) categorized by their KRAS genotype status (*x* axis). Gray: wildtype KRAS, red: known gain-of-function variants. Middle: Growth in low attachment of HA1E cells (z-score, color bar), or GILA score, for each variant (columns, ordered as in Fig. 2d) at 7 and 14 days.. Bottom: Mutation prevalence (log_2_(counts+1) of variant occurrences) in the COSMIC database (top) and a pan-cancer curated set (bottom), for each variant. **c**. Gene programs association with variants. Top: Difference (dot color) in mean program score (top) or mean PC score (bottom) between a variant (columns) and WT overexpressing cells and the significance of this difference (dot size, -log_10_(P-value), Kolmogorov-Smirnov test, **Methods**) for each gene program (top, rows, by gene clustering genes, **Methods**), or each of the top 10 PCs (bottom, rows). Colored border: BH FDR<10%. **d,e**. Relation of variants to different clusters and cell cycle phases. Left: Proportion of cells (bar height) in each cell cluster (**d,** as in Extended Data Fig. 1g) or cell cycle phase (**e**) (rows) derived for each variant (columns), annotated at the top with significance from a chi-square test comparing the cell state distribution of each variant with that of WT overexpressing cells (-log_10_(p-value)).. Right: UMAP embedding of single cell profiles, colored by cell clusters (**d**) or cell cycle phase (**e**). **f**. Relation of variants to KRAS gene structure. sc-eVIP scores (*y* axis) of each variant (dot, colored by the variant class) and its position along the KRAS gene (*x* axis, annotated by domain). **g**. Variant induced shift in cell distributions. Density map of cell profiles in a UMAP embedding, comparing the density of cells overexpressing variants in each of 3 classes to either the WT TP53 allele (grey, top) or unassigned cells (purple, bottom). **h**. UMAP embedding of single cell profiles colored by representative PCs.

**Extended Data Figure 9.**
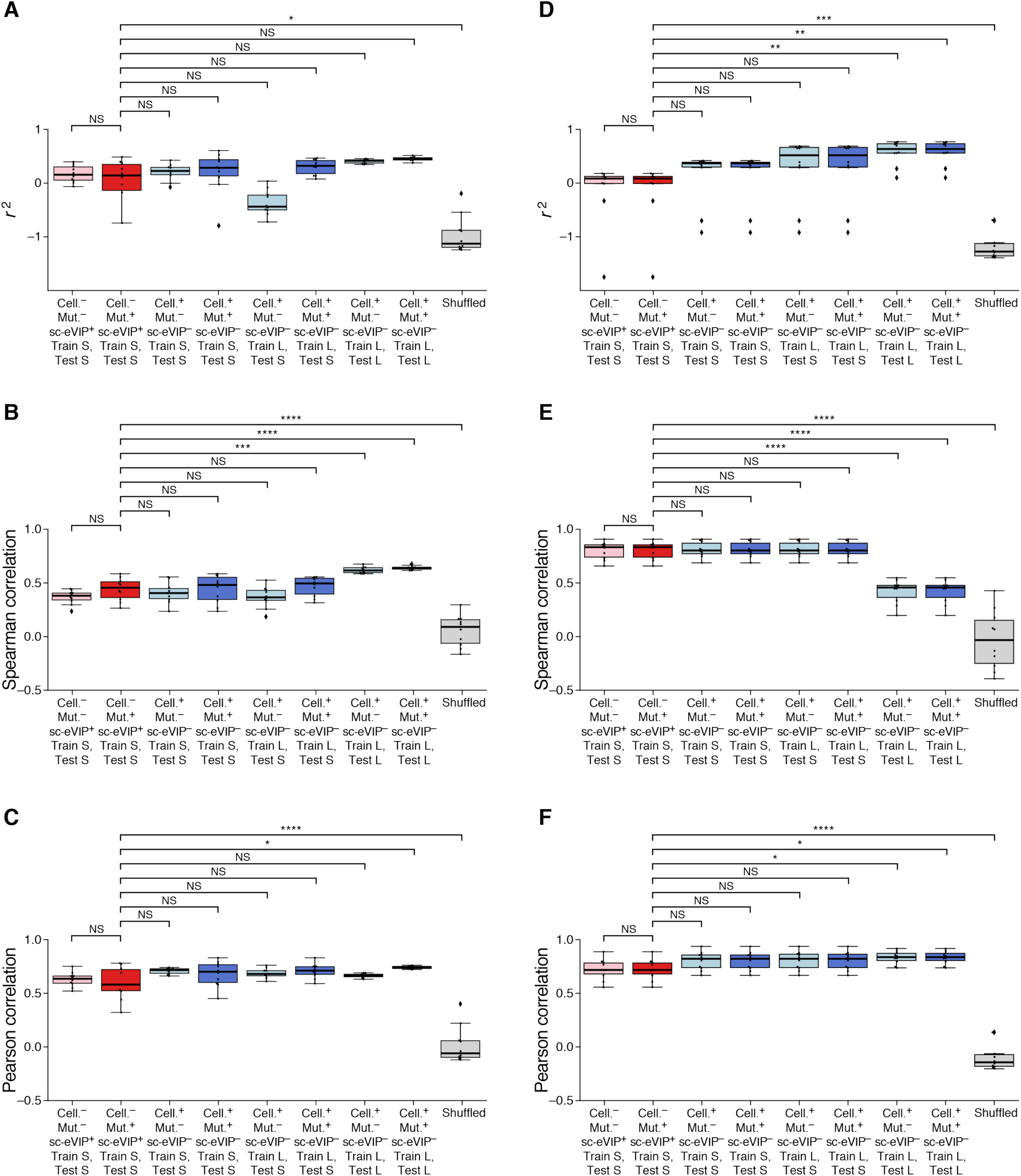
Performance of models predicting mutation prevalence for TP53 and KRAS variants. For each of TP53 (**a-c**) and KRAS variants (**d-f**), shown is the performance of different generalized models (*a* axis) by either *r*^2^ (**a,d** *y* axis, relative to a model that predicts the average mutation prevalence), or either Spearman (**b,e**) or Pearson (**c,f**) correlation coefficient between mutation prevalence and predictions. S: subset of variants profiled in this study; L: large dataset consisting of thousands of TP53 variants or tens of KRAS variants. Cell: functional assays data; Mut: mutational signature data. Shuffled: model which shuffles the observed variant prevalence across variants. Boxes are colored by use of mutational signatures (dark), sc-eVIP scores (red), and functional assays (blue). Boxplot shows the median, and its ends represent the 25% and 75% quartiles, with whiskers extending between (25% quartile - 1.5 interquartile range) and (75% quartile + 1.5 interquartile range) or the most extreme values in the data, if they fall within this range.

**Supplementary Table 1. Properties of TP53 variants.** The columns represent the name of the variant (Variant), its position in the amino acid sequence (Position), the original base(s) in the ORF (From), the base(s) the variant produces (To), whether the variant involves a single or multiple base change (Mutation type), whether the variant is a control synonymous, ExAC or unknown (Control status), whether the variant passed quality control and is in the library (Library synthesis), the number of cells per variant (Cells/variant), the average expression of the variant barcode in UMIs per 10,000 UMIs (Variant expression), Hotelling’s T^2^ statistic representing the sc-eVIP score (HotellingT2), the FDR (FDR.HotellingT2), the functional class assigned to the variant (Variant functional class), the variant prevalence in the pan-cancer dataset (Count(pancan)) and the variant prevalence in ExAC (Count (ExAC)).

**Supplementary Table 2. Properties of KRAS variants.** The columns represent the name of the variant (Variant), its position in the amino acid sequence (Position), the original base(s) in the ORF (From), the base(s) the variant produces (To), whether the variant involves a single or multiple base change (Mutation type), whether the variant is a control synonymous, ExAC or unknown (Control status), whether the variant passed quality control and is in the library (Library synthesis), the number of cells per variant (Cells/variant), the average expression of the variant barcode in UMIs per 10,000 UMIs (Variant expression), Hotelling’s T^2^ statistic representing the sc-eVIP score (HotellingT2), the FDR (FDR.HotellingT2), the functional class assigned to the variant (Variant functional class), the variant prevalence in the pan-cancer dataset (Count(pancan)) and the variant prevalence in ExAC (Count (ExAC)).

## REFERENCES

AACR Project GENIE Consortium (2017). AACR Project GENIE: Powering Precision Medicine through an International Consortium. Cancer Discov. 7, 818–831.

Adamson, B., Norman, T.M., Jost, M., Cho, M.Y., Nuñez, J.K., Chen, Y., Villalta, J.E., Gilbert, L.A., Horlbeck, M.A., Hein, M.Y., et al. (2016). A Multiplexed Single-Cell CRISPR Screening Platform Enables Systematic Dissection of the Unfolded Protein Response. Cell 167, 1867–1882.e21.

Akagi, K., Uchibori, R., Yamaguchi, K., Kurosawa, K., Tanaka, Y., and Kozu, T. (2007). Characterization of a novel oncogenic K-ras mutation in colon cancer. Biochem. Biophys. Res. Commun. 352, 728–732.

Alexandrov, L.B., Nik-Zainal, S., Wedge, D.C., Aparicio, S.A.J.R., Behjati, S., Biankin, A.V., Bignell, G.R., Bolli, N., Borg, A., Børresen-Dale, A.-L., et al. (2013). Signatures of mutational processes in human cancer. Nature 500, 415–421.

Alexandrov, L.B., Jones, P.H., Wedge, D.C., Sale, J.E., Campbell, P.J., Nik-Zainal, S., and Stratton, M.R. (2015). Clock-like mutational processes in human somatic cells. Nat. Genet. 47, 1402–1407.

Anzalone, A.V., Randolph, P.B., Davis, J.R., Sousa, A.A., Koblan, L.W., Levy, J.M., Chen, P.J., Wilson, C., Newby, G.A., Raguram, A., et al. (2019). Search-and-replace genome editing without double-strand breaks or donor DNA. Nature 576, 149–157.

Bailey, M.H., Tokheim, C., Porta-Pardo, E., Sengupta, S., Bertrand, D., Weerasinghe, A., Colaprico, A., Wendl, M.C., Kim, J., Reardon, B., et al. (2018). Comprehensive Characterization of Cancer Driver Genes and Mutations. Cell 174, 1034–1035.

Benson, D.A., Cavanaugh, M., Clark, K., Karsch-Mizrachi, I., Lipman, D.J., Ostell, J., and Sayers, E.W. (2013). GenBank. Nucleic Acids Res. 41, D36–D42.

Berger, A.H., Brooks, A.N., Wu, X., Shrestha, Y., Chouinard, C., Piccioni, F., Bagul, M., Kamburov, A., Imielinski, M., Hogstrom, L., et al. (2016). High-throughput phenotyping of lung cancer somatic mutations. Cancer Cell 30, 214–228.

Blondel, V.D., Guillaume, J.-L., Lambiotte, R., and Lefebvre, E. (2008). Fast unfolding of communities in large networks.

Boettcher, S., Miller, P.G., Sharma, R., McConkey, M., Leventhal, M., Krivtsov, A.V., Giacomelli, A.O., Wong, W., Kim, J., Chao, S., et al. (2019). A dominant-negative effect drives selection of TP53 missense mutations in myeloid malignancies. Science 365, 599–604.

Brenan, L., Andreev, A., Cohen, O., Pantel, S., Kamburov, A., Cacchiarelli, D., Persky, N.S., Zhu, C., Bagul, M., Goetz, E.M., et al. (2016). Phenotypic Characterization of a Comprehensive Set of MAPK1/ERK2 Missense Mutants. Cell Rep. 17, 1171–1183.

Brodin, P., Jojic, V., Gao, T., Bhattacharya, S., Angel, C.J.L., Furman, D., Shen-Orr, S., Dekker, C.L., Swan, G.E., Butte, A.J., et al. (2015). Variation in the human immune system is largely driven by non-heritable influences. Cell 160, 37–47.

Buschmann, T., and Bystrykh, L.V. (2013). Levenshtein error-correcting barcodes for multiplexed DNA sequencing. BMC Bioinformatics 14, 272.

Cao, J., Packer, J.S., Ramani, V., Cusanovich, D.A., Huynh, C., Daza, R., Qiu, X., Lee, C., Furlan, S.N., Steemers, F.J., et al. (2017). Comprehensive single-cell transcriptional profiling of a multicellular organism. Science 357, 661–667.

Chang, M.T., Asthana, S., Gao, S.P., Lee, B.H., Chapman, J.S., Kandoth, C., Gao, J., Socci, N.D., Solit, D.B., Olshen, A.B., et al. (2016). Identifying recurrent mutations in cancer reveals widespread lineage diversity and mutational specificity. Nat. Biotechnol. 34, 155–163.

Cleary, B., Cong, L., Cheung, A., Lander, E.S., and Regev, A. (2017). Efficient Generation of Transcriptomic Profiles by Random Composite Measurements. Cell 171, 1424–1436.e18.

Datlinger, P., Rendeiro, A.F., Boenke, T., Krausgruber, T., Barreca, D., and Bock, C. (2019). Ultra-high throughput single-cell RNA sequencing by combinatorial fluidic indexing.

Dixit, A. (2016). Correcting Chimeric Crosstalk in Single Cell RNA-seq Experiments.

Dixit, A., Parnas, O., Li, B., Chen, J., Fulco, C.P., Jerby-Arnon, L., Marjanovic, N.D., Dionne, D., Burks, T., Raychowdhury, R., et al. (2016). Perturb-Seq: Dissecting Molecular Circuits with Scalable Single-Cell RNA Profiling of Pooled Genetic Screens. Cell 167, 1853–1866.e17.

Dogruluk, T., Tsang, Y.H., Espitia, M., Chen, F., Chen, T., Chong, Z., Appadurai, V., Dogruluk, A., Eterovic, A.K., Bonnen, P.E., et al. (2015). Identification of Variant-Specific Functions of PIK3CA by Rapid Phenotyping of Rare Mutations. Cancer Res. 75, 5341–5354.

Dubovik, T., Starosvetsky, E., LeRoy, B., Normand, R., Admon, Y., Alpert, A., Ofran, Y., G’Sell, M., and Shen-Orr, S.S. (2018). Architecture of a multi-cellular polygenic network governing immune homeostasis.

Feldman, D., Singh, A., Schmid-Burgk, J.L., Carlson, R.J., Mezger, A., Garrity, A.J., Zhang, F., and Blainey, P.C. (2019). Optical Pooled Screens in Human Cells. Cell 179, 787–799.e17.

Figliuzzi, M., Jacquier, H., Schug, A., Tenaillon, O., and Weigt, M. (2016). Coevolutionary Landscape Inference and the Context-Dependence of Mutations in Beta-Lactamase TEM-1. Mol. Biol. Evol. 33, 268–280.

Findlay, G.M., Daza, R.M., Martin, B., Zhang, M.D., Leith, A.P., Gasperini, M., Janizek, J.D., Huang, X., Starita, L.M., and Shendure, J. (2018). Accurate classification of BRCA1 variants with saturation genome editing. Nature 562, 217–222.

Fischer, M. (2017). Census and evaluation of p53 target genes. Oncogene 36, 3943–3956.

Gao, Y., Chang, M.T., McKay, D., Na, N., Zhou, B., Yaeger, R., Torres, N.M., Muniz, K., Drosten, M., Barbacid, M., et al. (2018). Allele-Specific Mechanisms of Activation of MEK1 Mutants Determine Their Properties. Cancer Discov. 8, 648–661.

Gaudelli, N.M., Komor, A.C., Rees, H.A., Packer, M.S., Badran, A.H., Bryson, D.I., and Liu, D.R. (2017). Programmable base editing of A•T to G•C in genomic DNA without DNA cleavage. Nature 551, 464–471.

Giacomelli, A.O., Yang, X., Lintner, R.E., McFarland, J.M., Duby, M., Kim, J., Howard, T.P., Takeda, D.Y., Ly, S.H., Kim, E., et al. (2018). Mutational processes shape the landscape of TP53 mutations in human cancer. Nat. Genet. 50, 1381–1387.

Grau, J., Grosse, I., and Keilwagen, J. (2015). PRROC: computing and visualizing precision-recall and receiver operating characteristic curves in R. Bioinformatics 31, 2595–2597.

Hong, D.S., Fakih, M.G., Strickler, J.H., Desai, J., Durm, G.A., Shapiro, G.I., Falchook, G.S., Price, T.J., Sacher, A., Denlinger, C.S., et al. (2020). KRASG12C Inhibition with Sotorasib in Advanced Solid Tumors. N. Engl. J. Med. 383, 1207–1217.

Hopf, T.A., Ingraham, J.B., Poelwijk, F.J., Schärfe, C.P.I., Springer, M., Sander, C., and Marks, D.S. (2017). Mutation effects predicted from sequence co-variation. Nat. Biotechnol. 35, 128–135.

Hotelling, H. (1931). The Generalization of Student’s Ratio. The Annals of Mathematical Statistics 2, 360–378.

Jaitin, D.A., Weiner, A., Yofe, I., Lara-Astiaso, D., Keren-Shaul, H., David, E., Salame, T.M., Tanay, A., van Oudenaarden, A., and Amit, I. (2016). Dissecting Immune Circuits by Linking CRISPR-Pooled Screens with Single-Cell RNA-Seq. Cell 167, 1883–1896.e15.

Jeay, S., Gaulis, S., Ferretti, S., Bitter, H., Ito, M., Valat, T., Murakami, M., Ruetz, S., Guthy, D.A., Rynn, C., et al. (2015). A distinct p53 target gene set predicts for response to the selective p53–HDM2 inhibitor NVP-CGM097. eLife 4.

Kamburov, A., Lawrence, M.S., Polak, P., Leshchiner, I., Lage, K., Golub, T.R., Lander, E.S., and Getz, G. (2015). Comprehensive assessment of cancer missense mutation clustering in protein structures. Proc. Natl. Acad. Sci. U. S. A. 112, E5486–E5495.

Kim, E., Ilic, N., Shrestha, Y., Zou, L., Kamburov, A., Zhu, C., Yang, X., Lubonja, R., Tran, N., Nguyen, C., et al. (2016). Systematic Functional Interrogation of Rare Cancer Variants Identifies Oncogenic Alleles. Cancer Discov. 6, 714–726.

Kinker, G.S., Greenwald, A.C., Tal, R., Orlova, Z., Cuoco, M.S., McFarland, J.M., Warren, A., Rodman, C., Roth, J.A., Bender, S.A., et al. (2019). Pan-cancer single cell RNA-seq uncovers recurring programs of cellular heterogeneity.

Komor, A.C., Kim, Y.B., Packer, M.S., Zuris, J.A., and Liu, D.R. (2016). Programmable editing of a target base in genomic DNA without double-stranded DNA cleavage. Nature 533, 420–424.

Kotler, E., Shani, O., Goldfeld, G., Lotan-Pompan, M., Tarcic, O., Gershoni, A., Hopf, T.A., Marks, D.S., Oren, M., and Segal, E. (2018). A Systematic p53 Mutation Library Links Differential Functional Impact to Cancer Mutation Pattern and Evolutionary Conservation. Mol. Cell 71, 873.

Langmead, B., Trapnell, C., Pop, M., and Salzberg, S.L. (2009). Ultrafast and memory-efficient alignment of short DNA sequences to the human genome. Genome Biol. 10, R25.

Lawrence, M.S., Stojanov, P., Polak, P., Kryukov, G.V., Cibulskis, K., Sivachenko, A., Carter, S.L., Stewart, C., Mermel, C.H., Roberts, S.A., et al. (2013). Mutational heterogeneity in cancer and the search for new cancer-associated genes. Nature 499, 214–218.

Lawrence, M.S., Stojanov, P., Mermel, C.H., Robinson, J.T., Garraway, L.A., Golub, T.R., Meyerson, M., Gabriel, S.B., Lander, E.S., and Getz, G. (2014). Discovery and saturation analysis of cancer genes across 21 tumour types. Nature 505, 495–501.

Lebrigand, K., Magnone, V., Barbry, P., and Waldmann, R. (2020). High throughput error corrected Nanopore single cell transcriptome sequencing. Nat. Commun. 11, 4025.

Lek, M., Karczewski, K.J., Minikel, E.V., Samocha, K.E., Banks, E., Fennell, T., O’Donnell-Luria, A.H., Ware, J.S., Hill, A.J., Cummings, B.B., et al. (2016). Analysis of protein-coding genetic variation in 60,706 humans. Nature 536, 285–291.

Levine, J.H., Simonds, E.F., Bendall, S.C., Davis, K.L., Amir, E.-A.D., Tadmor, M.D., Litvin, O., Fienberg, H.G., Jager, A., Zunder, E.R., et al. (2015). Data-Driven Phenotypic Dissection of AML Reveals Progenitor-like Cells that Correlate with Prognosis. Cell 162, 184–197.

Li, Z., Del-Aguila, J.L., Dube, U., Budde, J., Martinez, R., Black, K., Xiao, Q., Cairns, N.J., Dominantly Inherited Alzheimer Network (DIAN), Dougherty, J.D., et al. (2018). Genetic variants associated with Alzheimer’s disease confer different cerebral cortex cell-type population structure. Genome Med. 10, 43.

Lu, J., Bera, A.K., Gondi, S., and Westover, K.D. (2018). KRAS Switch Mutants D33E and A59G Crystallize in the State 1 Conformation. Biochemistry 57, 324–333.

Ly, S.H. (2018). Investigation of KRAS Dependency Bypass and Functional Characterization of All Possible KRAS Missense Variants.

Ma, S., Zhang, B., LaFave, L.M., Earl, A.S., Chiang, Z., Hu, Y., Ding, J., Brack, A., Kartha, V.K., Tay, T., et al. (2020). Chromatin Potential Identified by Shared Single-Cell Profiling of RNA and Chromatin. Cell.

Macosko, E.Z., Basu, A., Satija, R., Nemesh, J., Shekhar, K., Goldman, M., Tirosh, I., Bialas, A.R., Kamitaki, N., Martersteck, E.M., et al. (2015). Highly Parallel Genome-wide Expression Profiling of Individual Cells Using Nanoliter Droplets. Cell 161, 1202–1214.

McFarland, J.M., Paolella, B.R., Warren, A., Geiger-Schuller, K., Shibue, T., Rothberg, M., Kuksenko, O., Colgan, W.N., Jones, A., Chambers, E., et al. (2020). Multiplexed single-cell transcriptional response profiling to define cancer vulnerabilities and therapeutic mechanism of action. Nat. Commun. 11, 4296.

Meyers, R.M., Bryan, J.G., McFarland, J.M., Weir, B.A., Sizemore, A.E., Xu, H., Dharia, N.V., Montgomery, P.G., Cowley, G.S., Pantel, S., et al. (2017). Computational correction of copy number effect improves specificity of CRISPR-Cas9 essentiality screens in cancer cells. Nat. Genet. 49, 1779– 1784.

Noble, W.S. (2009). How does multiple testing correction work? Nature Biotechnology 27, 1135–1137.

Rehm, H.L., and Fowler, D.M. (2019). Keeping up with the genomes: scaling genomic variant interpretation. Genome Med. 12, 5.

Rodriguez, J.M., Rodriguez-Rivas, J., Di Domenico, T., Vázquez, J., Valencia, A., and Tress, M.L. (2018). APPRIS 2017: principal isoforms for multiple gene sets. Nucleic Acids Res. 46, D213–D217.

Rohban, M.H., Singh, S., Wu, X., Berthet, J.B., Bray, M.-A., Shrestha, Y., Varelas, X., Boehm, J.S., and Carpenter, A.E. (2017). Systematic morphological profiling of human gene and allele function via Cell Painting. Elife 6.

Rosenberg, A.B., Roco, C.M., Muscat, R.A., Kuchina, A., Sample, P., Yao, Z., Graybuck, L.T., Peeler, D.J., Mukherjee, S., Chen, W., et al. (2018). Single-cell profiling of the developing mouse brain and spinal cord with split-pool barcoding. Science 360, 176–182.

Rotem, A., Janzer, A., Izar, B., Ji, Z., Doench, J.G., Garraway, L.A., and Struhl, K. (2015). Alternative to the soft-agar assay that permits high-throughput drug and genetic screens for cellular transformation. Proc. Natl. Acad. Sci. U. S. A. 112, 5708–5713.

Sidore, A.M., Plesa, C., Samson, J.A., and Kosuri, S. (2019). DropSynth 2.0: high-fidelity multiplexed gene synthesis in emulsions.

Singh, A., Greninger, P., Rhodes, D., Koopman, L., Violette, S., Bardeesy, N., and Settleman, J. (2009). A gene expression signature associated with “K-Ras addiction” reveals regulators of EMT and tumor cell survival. Cancer Cell 15, 489–500.

Tate, J.G., Bamford, S., Jubb, H.C., Sondka, Z., Beare, D.M., Bindal, N., Boutselakis, H., Cole, C.G., Creatore, C., Dawson, E., et al. (2019). COSMIC: the Catalogue Of Somatic Mutations In Cancer. Nucleic Acids Res. 47, D941–D947.

UniProt Consortium (2019). UniProt: a worldwide hub of protein knowledge. Nucleic Acids Res. 47, D506–D515.

Volden, R., and Vollmers, C. (2020). Highly Multiplexed Single-Cell Full-Length cDNA Sequencing of human immune cells with 10X Genomics and R2C2.

Wolf, F.A., Angerer, P., and Theis, F.J. (2018). SCANPY: large-scale single-cell gene expression data analysis. Genome Biol. 19, 15.

Yu, K., Lin, C.-C.J., Hatcher, A., Lozzi, B., Kong, K., Huang-Hobbs, E., Cheng, Y.-T., Beechar, V.B., Zhu, W., Zhang, Y., et al. (2020). PIK3CA variants selectively initiate brain hyperactivity during gliomagenesis. Nature 578, 166–171.

Zehir, A., Benayed, R., Shah, R.H., Syed, A., Middha, S., Kim, H.R., Srinivasan, P., Gao, J., Chakravarty, D., Devlin, S.M., et al. (2017). Mutational landscape of metastatic cancer revealed from prospective clinical sequencing of 10,000 patients. Nat. Med. 23, 703–713.

Zheng, G.X.Y., Terry, J.M., Belgrader, P., Ryvkin, P., Bent, Z.W., Wilson, R., Ziraldo, S.B., Wheeler, T.D., McDermott, G.P., Zhu, J., et al. (2017). Massively parallel digital transcriptional profiling of single cells. Nat. Commun. 8, 14049. FoundationOne CDx.

